# High-content profiling reveals a unified model of copper ionophore dependent cell death in oesophageal adenocarcinoma

**DOI:** 10.1101/2021.10.05.463189

**Authors:** Rebecca E. Hughes, Richard J. R. Elliott, Xiaodun Li, Alison F. Munro, Ashraff Makda, Roderick N. Carter, Nicholas M. Morton, Kenji Fujihara, Nicholas J. Clemons, Rebecca Fitzgerald, J. Robert O’Neill, Ted Hupp, Neil O. Carragher

## Abstract

**Background and Aims:** Oesophageal adenocarcinoma (OAC) is of increasing global concern due to increasing incidence, a lack of effective treatments, and poor prognosis. Therapeutic target discovery and clinical trials have been hindered by the heterogeneity of the disease, lack of driver mutations, and the dominance of large-scale genomic rearrangements. In this work we have characterised three potent and selective hit compounds identified in an innovative high-content phenotypic screening assay. The three hits include two approved drugs; elesclomol and disulfiram, and another small molecule compound, ammonium pyrrolidinedithiocarbamate. We uncover their mechanism of action, discover a targetable vulnerability, and gain insight into drug sensitivity for biomarker-based clinical trials in OAC.

**Methods:** Elesclomol, disulfiram, and ammonium pyrrolidinedithiocarbamate were systematically characterised across panels of oesophageal cell lines and patient-derived organoids. Drug treated oesophageal cell lines were morphologically profiled using a high-content, imaging platform. Compounds were assessed for efficacy across patient-derived organoids. Metabolomics and transcriptomics were assessed for the identification of oesophageal-cancer specific drug mechanisms and patient stratification hypotheses.

**Results:** High-content profiling revealed that all three compounds were highly selective for OAC over tissue-matched controls. Comparison of gene expression and morphological signatures unveiled a unified mechanism of action involving the accumulation of copper selectively in cancer cells, leading to dysregulation of proteostasis and cancer cell death. Basal omic analyses revealed proteasome and metabolic markers of drug sensitivity, forming the basis for biomarker-based clinical trials in OAC.

**Conclusions:** Integrated analysis of high-content imaging, transcriptomic and metabolomic data has revealed a new therapeutic mechanism for the treatment of OAC and represents an alternative target-agnostic drug discovery strategy.

## Introduction

Oesophageal cancer is an increasing global concern due to increasing incidence, a lack of effective treatments, and poor prognosis ^1^. Combined, the two major histological subtypes; oesophageal adenocarcinoma (OAC) and oesophageal squamous cell carcinoma (OSCC), represent the sixth leading cause of cancer deaths worldwide with less than one in five patients surviving five years from diagnosis ^2^. A shift in epidemiology over the last 50 years has meant the incidence of OAC now vastly exceeds that of OSCC in western countries ^1^, accounting for more than 90 % of oesophageal cancers in the United States ^3^.

Whole genome sequencing of clinical samples has recently highlighted OAC as highly heterogeneous, characterized by frequent large scale genomic rearrangements and copy number alterations ^4^. Despite recent advances in targeted therapies for tumours expressing HER2, VEGFR2 or PDL1 ^5–9^, survival rates remain low for a large proportion of patients and overall 5-year survival rates remain less than 20 %. This highlights the limitations of modern target-based drug discovery strategies to impact upon complex heterogeneous diseases such as OAC. Phenotypic drug discovery, a complementary approach to target-based drug discovery ^10,11^, describes the screening and selection of compounds based on quantifiable phenotypic endpoints without prior knowledge of the drug target. It is therefore an attractive strategy for heterogeneous diseases where there is a lack of understanding of disease biology and actionable targets.

This current study focuses on the characterisation of three small molecule compounds; elesclomol, disulfiram, and ammonium pyrrolidinedithiocarbamate, which show remarkable potency and selectivity for OAC over tissue-matched controls. These three molecules were identified using an image-based phenotypic screening assay we have previously described ^12^, using the high-content Cell Painting assay ^13,14^ to quantify the morphological response to a total of 19,555 small molecules. (Supplementary Table 1) across a panel of genetically distinct human OAC cell lines.

In this study, we report for the first time the full list of primary screening hits (Supplementary Table 1) and dose response hit validation studies (Supplementary Table 6) from target annotated LOPAC and Prestwick FDA approved compounds. The latest advances in multiparametric image-based high-content profiling were then employed to determine the activity of the most potent and selective hit molecules across panels of oesophageal cell lines and patient-derived organoids. We then systematically explored potential mechanisms of drug action of two hit compounds (elesclomol and disulfiram) which have already been approved for clinical use in a holistic approach to understand the molecular mechanisms which confer drug sensitivity and potentially identify new therapeutic targets and classes of small-molecules for the treatment of OAC.

Finally, transcriptomic and metabolomics analyses were utilised to determine factors affecting drug sensitivity generating patient stratification hypotheses for further validation. We believe the strategies we have employed in this study to identify and characterise highly selective hit compounds for OAC embraces the complexity of heterogeneous diseases and can be applied early on in the drug discovery pipeline to have a beneficial impact upon drug discovery success rates as a whole.

## Materials and Methods

### Cell culture

OAC lines were grown in RPMI-1640 (Life Technologies; #11875101) with 10% fetal bovine serum (FBS) (Life Technologies; #16140071) and 2mM L-glutamine (Life Technologies; #A2916801). CP-A and EPC2-hTERT cells were grown in KSFM (Life Technologies; #17005075) with 5g/L human recombinant epidermal growth factor and 50mg/L bovine pituitary extract. Supplementary Table 2 for cell line details.

Primary organoid cultures were derived from normal gastric and OAC cases included in the oesophageal cancer clinical and molecular stratification (OCCAMS)/international cancer genome consortium (ICGC) sequencing study. Detailed organoid culture and derivation method have been previously described in detail ^15^. Cells were seeded in complete medium then treated with elesclomol in 9-point half-log serial dilution for 6 days (maximal concentration 10 μM). Treatments were performed in technical duplicates and at least two biological replicates. Cell viability was assessed using CellTiter-Glo (Promega).

### Compound screening

Cells were seeded at 800-1500 cells per well in 50 μL into 384-well microplates (Greiner, #781091), for 24 hrs. Compounds were then added as 8 point semi log dose responses from 10μM before being incubated for a further 48 hrs. The Cell Painting protocol which uses multiplexed fluorescent dyes to visualise cellular and subcellular organelle and cytoskeletal morphology ^13,14^. Cells were fixed in 4% formaldehyde before permeabilisation in 0.1% Triton-X199. Finally the staining solution was added in 1% bovine serum albumin and incubated for 30 mins. (Supplementary Table 3). Four fields of view were captured per well using a 20x objective and five filters (Supplementary Table 3)

### Transcriptomic analyses

Cells were seeded at 8 x 10^4^ cells in 6-well plates for 24 hrs. Media was replaced with fresh media containing compound treatments (DMSO (0.1%), disulfiram (600 nM), or elesclomol (200 nM)) before further incubation for 6 hrs. Media was then removed and plates were washed twice with ice cold PBS before being snap frozen at −80. Cells were scraped and lysed using QIAshredders (#79654, QIAGEN) and Qiagen RNeasy Mini kit (#74104, QIAGEN) (with β-mercaptoethanol) according to manufacturer instructions, and included a DNase digestion step (#79254, QIAGEN). 100□ng of each sample was loaded into the NanoString nCounter Analysis System with the Human PanCancer Pathways and Metabolic Pathways panels. Raw counts were normalized to the internal positive controls and housekeeping genes using the nSolver 4.0 software. nSolver 4.0 software and the NanoStringDiff algorithm ^16^ was used for differential expression analysis. P-values were adjusted using the Benjamini-Yekutieli approach ^17^. Treatment induced analysis was carried out between the control (DMSO) and treatment (200 nM elesclomol) samples for the two sensitive cell lines (OAC-P4C and SK-GT-4) pooled together. N=3. Difference in cell lines was taken into account as a confounder in the analysis. Sensitive versus resistant OAC analysis was carried out between the DMSO treated sensitive cell lines (OAC-P4C, SK-GT-4, OE19, and ESO26) and the resistant cell lines (FLO-1, JH-EsoAd1, and ESO51). N=3.

### Cell metabolic assays

Cells were seeded at 8,000-10,000 cells per well in V7 Seahorse 24-well plates (250 μL per well) and incubated overnight. For treatment induced oxygen consumption rate experiments DMSO (0.1%) or elesclomol (50 and 200 nM) were added and incubated for 6 hrs. Compound incubated plates and basal plates were then washed with assay media (Seahorse XF DMEM assay media kit (Agilent; #103575-100), 10 mM glucose, 1 mM pyruvate, and 2 mM glutamax) twice before adding assay media to a final volume of 525 μL and incubated at 37 °C in CO_2_ free incubator for 30 minutes. Assays were run according to manufacturer’s instructions and cell count was used for normalisation. Supplementary Table 4 for cell line specific details.

### Intracellular copper quantification

5 x 10^6^ cells were seeded in a T175 flask and incubated for 24 hrs. Media was replaced with compound treatments (DMSO (0.1%), disulfiram (600 nM), or elesclomol (200 nM)) before further incubation for 6 hrs. Cells were washed in PBS, typsinised and counted. For each sample 2 x 10^6^ cells were pelleted and frozen at −80 °C. For analysis, samples were thawed and concentrated nitric acid was added (100 μL) and mixed. Samples were then vortexed and sonicated and left overnight at room temperature. Samples were made up to 1 mL using water and then further diluted fivefold prior to analysis of copper content by inductively coupled plasma mass spectrometry (ICP-MS).

## Results

### Elesclomol, disulfiram, and ammonium pyrrolidinedithiocarbamate are highly selectively cytotoxic towards OAC

We performed image-based high-content phenotypic profiling of 74 compound hits identified from a phenotypic screen (Supplementary Table 5 and 6) as dose responses across a panel of heterogeneous oesophageal cell lines to quantify both cell growth and morphometric response to compound treatments. Using such multivariate compound profiles we can gain a deeper understanding of compound activity, mechanism of action, and selectivity ^18^. Across a panel of six OAC cell lines elesclomol had the greatest potency in the low nanomolar range, followed by disulfiram and then ammonium pyrrolidinedithiocarbamate (Figure 1A and Supplementary Table 7). Critically the dose responses show no toxicity in either of the tissue-matched non-transformed control cell lines (CP-A a Barrett’s oesophagus cell line and EPC2-hTERT a squamous oesophageal cell line) and both continued to proliferate at their normal rate (Figure 1A, Supplementary Figure 1 for structures).

**Figure 1:**
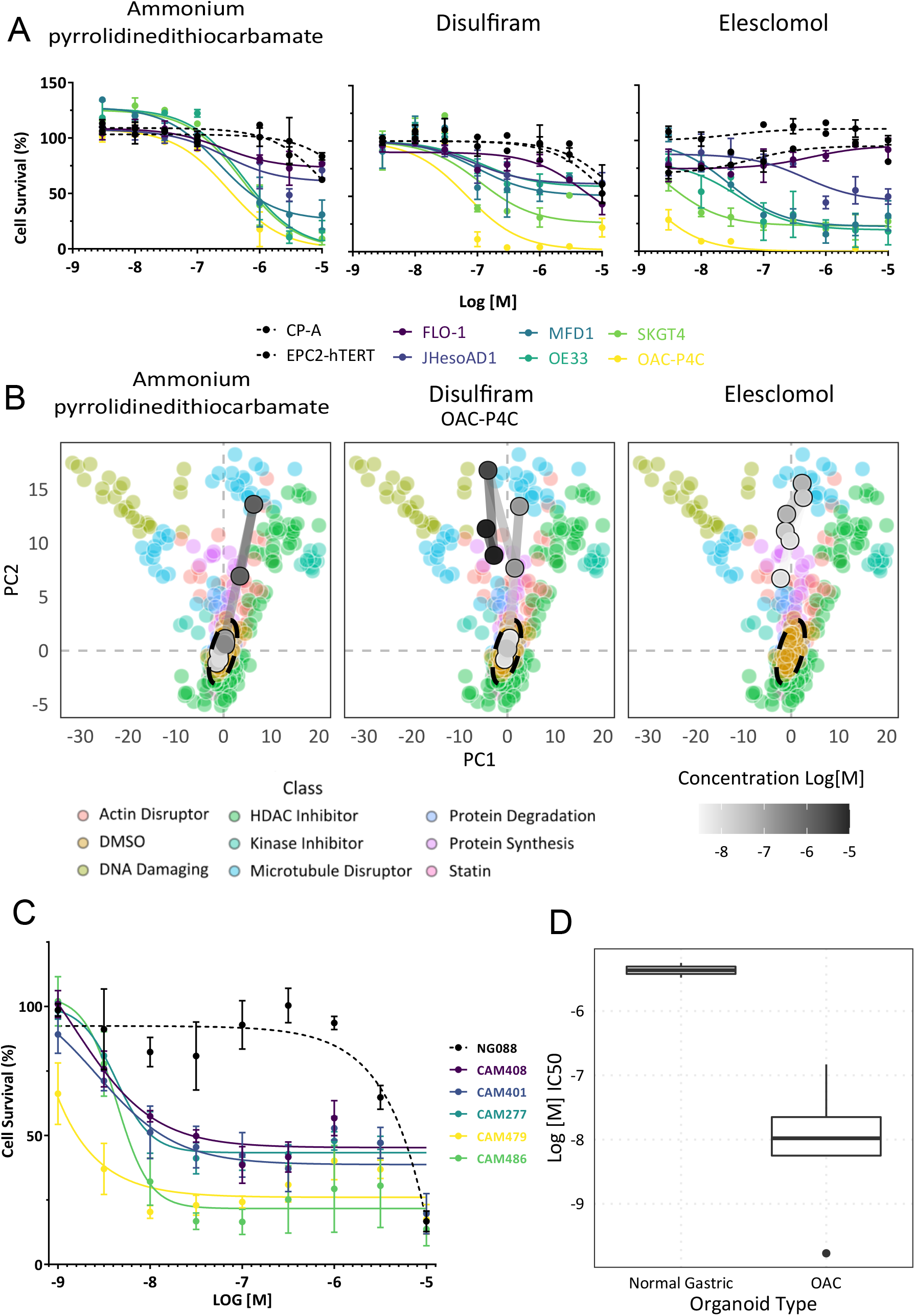
Dose responses for ammonium pyrrolidinedithiocarbamate, disulfiram and elesclomol. A) Univariate validation dose responses across cell panel. B) First two principal components of multivariate phenotypic dose responses in OAC-P4C (most sensitive cell line) overlaid on the reference library of compounds. C) Patient derived organoid dose responses for elesclomol. NG088 - normal gastric control organoid. D) Comparison of IC_50_ for normal gastric and OAC organoids, p < 0.005.

To test the most promising of the three compounds in patient-derived organoids, one gastric epithelial and five OAC patient-derived organoids were treated with elesclomol (Figure 2). Elesclomol showed a similar potency in the organoids compared to the adherent OAC cell lines, with IC_50_ values in the low nanomolar range (Supplementary Table 8). Elesclomol was also highly selective, inhibiting the viability of tumour derived organoids and not the normal gastric organoid NG088 (Figure 1C and D). These data demonstrate that elesclomol, disulfiram, and ammonium pyrrolidinedithiocarbamate are highly potent and selective for OAC over tissue-matched control cells in both 2D and 3D models.

**Figure 2:**
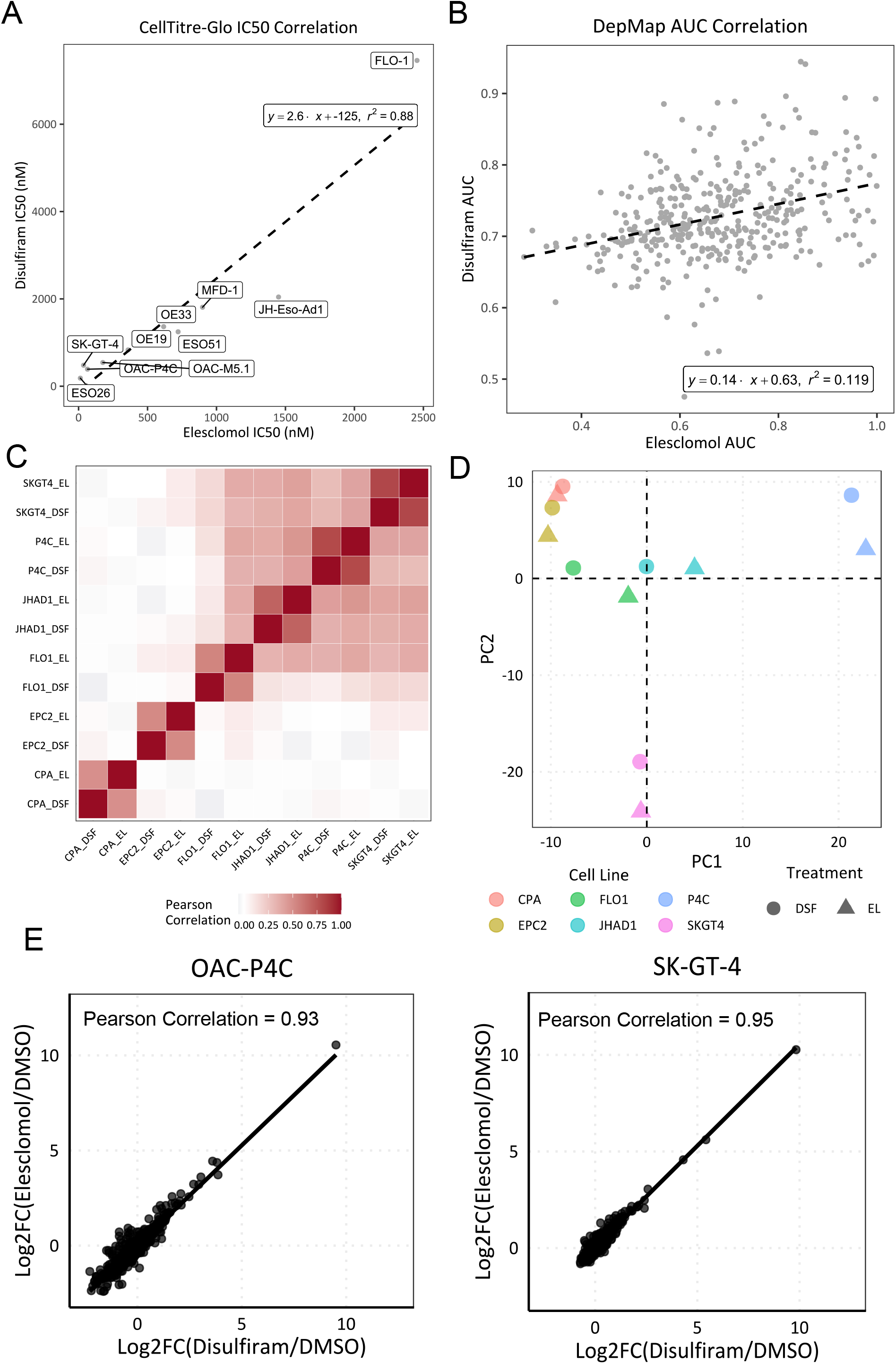
Correlations between disulfiram and elesclomol. A) IC_50_s across expanded cell panel. B) Pan cancer correlation of area under curve. Data from DepMap PRISM Repurposing Secondary Screen 19Q4, n=373. C) Correlation heatmap of log2 fold gene expression changes across both OAC and tissue-matched controls. D) First two principal components of average induced gene expression changes coloured by cell line. E) Correlation of gene expression changes in individual OAC cell lines. n = 3. DSF - Disulfiram, EL - Elesclomol.

Using the phenotypic information from the concentration responses we built multivariate dose responses, tracking cellular morphological changes as a product of compound concentration. We have shown previously, using a reference library of well-annotated compounds (Supplementary Table 9) that compounds with a shared mechanism of action cluster by induced cellular phenotype ^12^. The phenotypic dose responses show that both disulfiram and ammonium pyrrolidinedithiocarbamate move from phenotypically inactive at low concentrations, clustering with the DMSO, to phenotypically active at higher concentrations (Figure 1B), while elesclomol is active at all concentrations tested. All three compounds cluster away from the known pharmacological classes in the reference library, suggesting a mechanism of action that is distinct from the reference library (Figure 1B). Correlation of our multivariate phenotypic dose response data shows all three compounds induce a shared morphological response (Pearson correlation 0.61 and 0.81 within OAC-P4C and SK-GT-4 cells respectively) (Supplementary Figure 2 for images), suggesting a potentially shared mechanism between the compounds. Of note, the compounds do not cause any phenotypic changes in the CP-A or the EPC2-hTERT cell lines across the dose responses (Supplementary Figure 3), further demonstrating strong selectivity.

### Elesclomol and disulfiram act through a unified mechanism of action in OAC cells

Previous studies have indicated that these three compounds have varied targets and therefore likely act via multiple distinct mechanisms ^19–25^. However, our data indicates that all three compounds induce a similar morphological signature from multivariate phenotypic analysis and also share the same sensitivity profile based on our univariate viability assessment across the panel of cell lines with OAC-P4C showing the strongest responses and FLO-1 showing the weakest (Figure 1). We quantified the sensitivity profile by Pearson correlation of IC_50_ values for disulfiram and elesclomol across an expanded panel of OAC cell lines which confirmed a shared sensitivity profile across OAC cell lines (Pearson 0.94, p-value <0.001) (Figure 2A). Similar analyses using data across 373 cancer cell lines from multiple tumour types from the Cancer Dependency Map (DepMap) ^26,27^ also demonstrated a significant correlation (Pearson 0.34, p-value <0.001) (Figure 2B). Combined, these data suggest that the compounds act through a shared mechanism in OAC.

We explored the mechanism further using the two most potent compounds which are clinically approved drugs; elesclomol and disulfiram, although we believe the mechanism is shared across all three compounds.

To further confirm a shared mechanism we used compound induced gene expression signatures. Comparison of signatures can be used to discover new connections among compounds as well as to identify targets and mechanisms of action in a more complex setting than single gene contributions ^28,29^. We used the NanoString nCounter platform to quantify and compare transcript expression following treatment with elesclomol or disulfiram. A correlation heatmap of cell lines and treatments showed that the compound induced signatures are very similar within a given cell line and to a lesser extent across cell lines (Figure 2C). Secondly, the OAC lines formed a distinct cluster to the tissue-matched control lines (CP-A and EPC2-hTERT) (Figure 2C). Pearson correlation and principal component analysis showed there was a very strong relationship between the two treatments within a cell line (Figure 2C and D and Supplementary Figure 4), demonstrating that disulfiram and elesclomol induce the same gene expression signature. Interestingly, neither elesclomol nor disulfiram induced any significant gene expression changes in the two tissue-matched controls (Supplementary Figure 5).

### Elucidation of mechanisms conferring sensitivity to disulfiram and elesclomol

Given the clear variability in OAC cell line sensitivity to disulfiram and elesclomol, we sought to identify the mechanisms which confer sensitivity through the integration of basal transcriptomic and metabolomics data.

Using both a differential expression analysis of nine OAC lines profiled with the NanoString nCounter platform, and a pan cancer correlation of gene expression using the Genomics of Drug Sensitivity in Cancer data ^30–32^ we identified a number of potential biomarkers of sensitivity, including methionine sulfoxide reductase B2 (MSRB2) and B3 (MSRB3), and glutathione peroxidase 8 (GPX8) (Figure 3A and B). We then applied gene set enrichment analysis (GSEA) to the Genomics of Drug Sensitivity data ^31^ for seven OAC cell lines to reveal significant associations at the biological pathway level. GSEA of hallmark gene sets revealed 10 that positively correlated and 11 that negatively correlated with IC_50_ (p < 0.05) (Figure 3C). MYC targets are the top two gene sets positively associated with elesclomol sensitivity, from which the top differentially expressed genes are ubiquitin enzymes and proteasome subunits (Supplementary Table 10). In concordance with this the unfolded protein response is the fifth gene set identified. Furthermore, GSEA of KEGG (Kyoto Encyclopedia of Genes and Genomes) gene sets, revealed 18 that were significantly enriched and positively correlated with elesclomol sensitivity, of which the proteasome was the top gene set (Figure 3D). Both analyses also identify bile acid metabolism and other aspects of metabolism, particularly lipid and fatty acid metabolism. Of note, cysteine and methionine metabolism was identified as the sixth most significant KEGG pathway from the GSEA analysis, giving further confidence to the identification of the methionine sulfoxide reductase genes at the single gene level.

**Figure 3:**
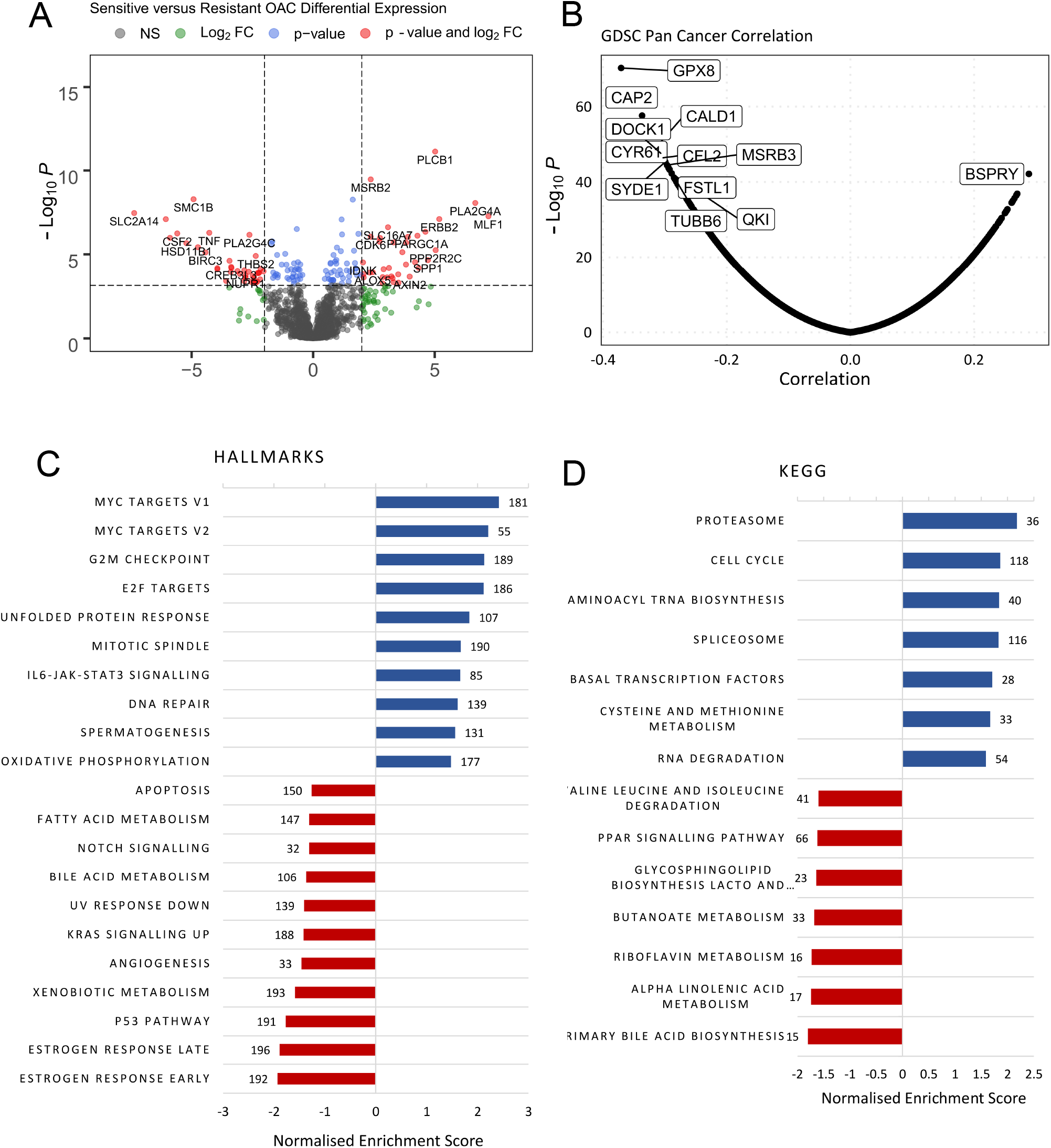
Sensitivity analysis. A) Differential expression analysis. Sensitive cell lines: OAC-P4C, SK-GT-4, OE19, ESO26. Resistant cell lines: FLO-1, JH-EsoAd1, ESO51. N=3. P value threshold equivalent to adjusted p value 0.05. Gene total = 1397. B) Correlation analyses of Genomics of Drug Sensitivity in Cancer. RMA normalised expression values for 17,737 genes against elesclomol area under the curve for 916 cell lines. C) GSEA Hallmarks analysis. Using the MSigDB Hallmark collection of 50 gene sets. D) GSEA KEGG analysis. Using the canonical pathways KEGG collection of 186 gene sets. Numbers represents gene set size. Pearson was used as a rank metric.

Using the Cancer Cell Line Encyclopedia (CCLE) metabolomics data ^33^ and MetaboAnalyst 5.0 we identified several aspects of metabolism to be associated with sensitivity including taurine and hypotaurin metabolism, bile acid biosynthesis, fatty acid metabolism, and mitochondrial beta-oxidation of long chain fatty acids (Figure 4 and Supplementary Figure 6). This corroborates the transcriptomic studies above, with both studies implicating bile acid and lipid metabolism pathways in compound sensitivity.

**Figure 4:**
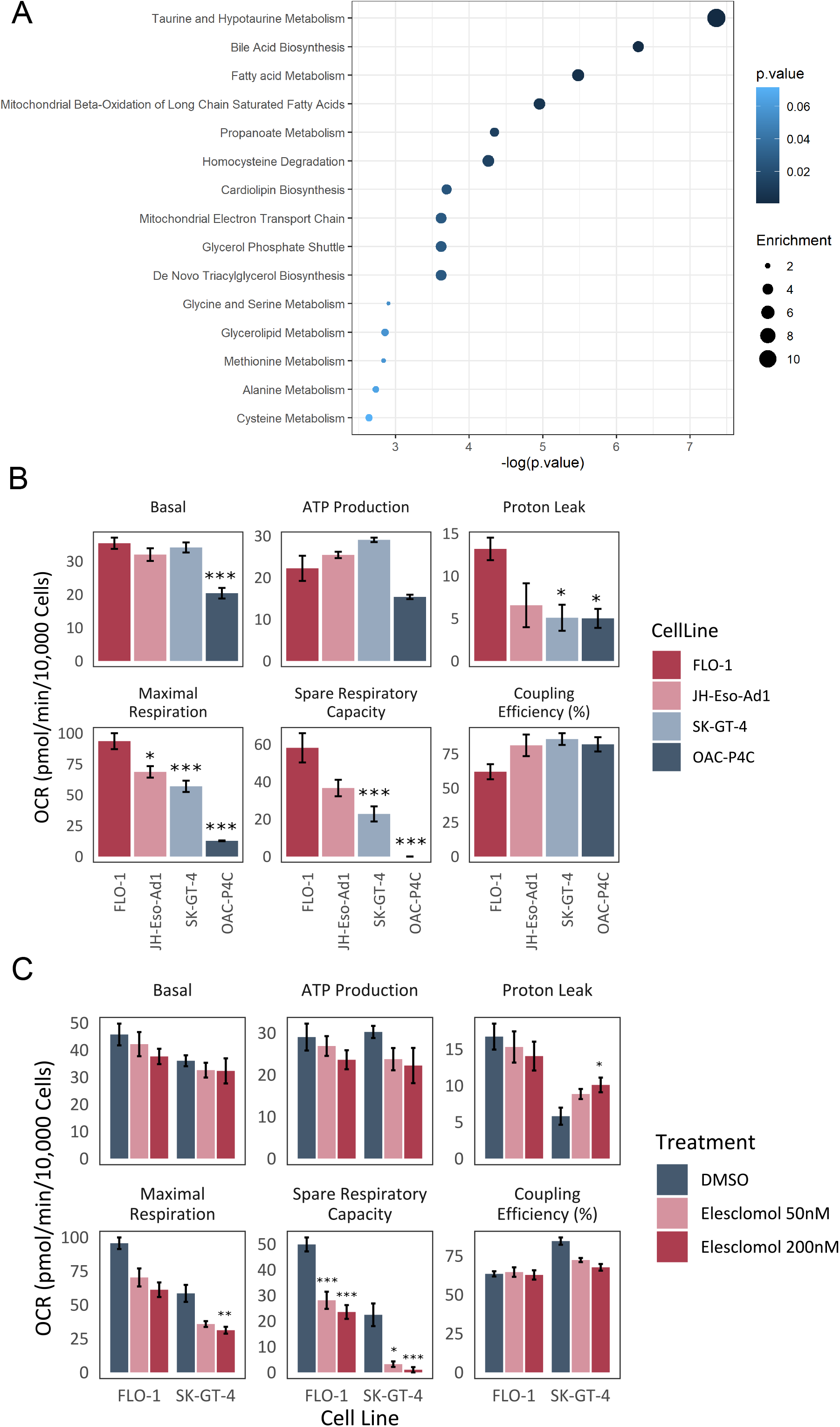
Metabolic effects. A) Quantitative enrichment analysis CCLE metabolomics data for elesclomol sensitivity. Metabolite sets ranked by significance. Point size represents enrichment value. B) Basal cell line seahorse parameters. Significance indicated when compared to most resistant cell line, FLO-1. C) Effect of elesclomol treatment on mitochondrial function. Exemplar OCR traces for a resistant and sensitive cell line when treated with elesclomol. B and C normalised to 10,000 cells, n=3. Error bars indicate SEM. 1-ANOVA and Tukeys post hoc test for significance. * P>0.05, ** p>0.01, *** p>0.005.

Investigation of OAC metabolism using the Seahorse Cell Mito Stress test revealed no significant difference in basal respiration between the sensitive and resistant OAC lines (Figure 4). However there are significant differences in the maximal respiration rate and the spare respiratory capacity, and both parameters are also significantly correlated with sensitivity, as determined by IC_50_ correlation (Kendal tau max respiration = 0.87, spare capacity = 0.89). This suggests energetic flexibility may play a role in sensitivity ^34^. Further investigation of the contribution of individual substrates, including fatty acids, to metabolism may reveal additional differences defining sensitivity.

Examining the effects of elesclomol drug treatment on respiratory function, we found that it reduced mitochondrial function in both sensitive and insensitive OAC cell lines, indicated by reduced maximal respiration and spare respiratory capacity, which was more significant in the sensitive cells (Figure 4), further implicating metabolism.

### Compound induced gene expression signatures reveals increased levels of misfolded proteins

In an unbiased approach to studying the mechanism of action of these compounds, we focused on the transcriptomic profile of elesclomol treatment in OAC. NanoString differential expression analysis ^16^ revealed strong induction of multiple heat shock and growth arrest and DNA damage genes, and heme oxygenase I (Figure 5A and B). We then used Ingenuity Pathway Analysis to define deregulated pathways in a compound exposure setting to elucidate mechanism of action. Pathway analysis revealed strong and consistent modulation of SAPK/JNK signalling, NRF2-mediated oxidative stress response, unfolded protein response, endoplasmic reticulum stress, and p53 signalling as the top five altered pathways after elesclomol treatment, known to arise from increased levels of misfolded proteins ^35^ (Figure 5C).

**Figure 5:**
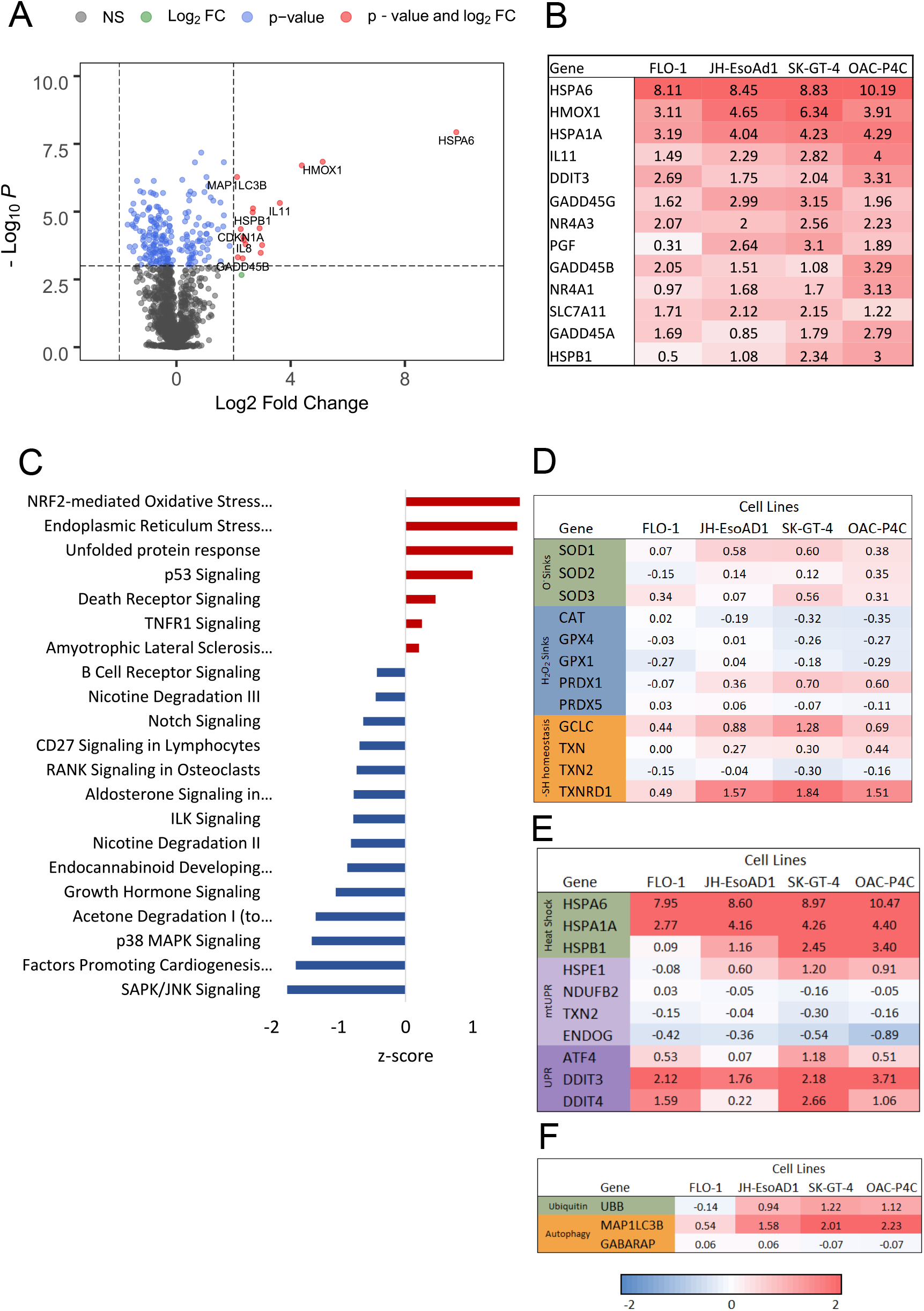
Elesclomol induced gene expression signature. A) Differential expression analysis for elesclomol treatment (200 nM for 6 h) vs DSMO in the two most sensitive cell lines OAC-P4C and SK-GT-4. P value cut off equivalent to adjusted p value 0.05. n = 3. B) Log2 fold change for top significant genes C) Top pathway Z-scores (by p-value) identified in ingenuity pathway analysis in the sensitive cell lines (OAC-P4C and SK-GT-4). D-F) Effect of elesclomol treatment (200 nM for 6 h) on the transcriptional profile of antioxidant enzymes (D), heat shock response and unfolded protein response (E), autophagy and proteasomal degradation (F). Log2 fold change values for gene expression across OAC cell lines using NanoString data. n=3

### Compound activity is not associated with reactive oxygen species

The proposed mechanism of elesclomol is induction of reactive oxygen species (ROS) ^36^. However, assessment of the enzymes involved in maintenance of redox homeostasis did not reveal any major induction of antioxidant enzymes in the elesclomol induced signature from OAC treated cells (Figure 3D). In fact, H_2_O_2_ sink enzymes glutathione peroxidase 1 and 4 (GPX1, GPX4), and catalase (CAT) were transcriptionally downregulated (Figure 5D).

Furthermore, if ROS levels were responsible for the activity of disulfiram and elesclomol in OAC, then we would expect the sensitivity of disulfiram and elesclomol to correlate with H_2_O_2_ sensitivity. However, we found that FLO-1 cells are significantly more sensitive to oxidative stress than either OAC-P4C cells or the tissue-matched control EPC2-hTERT (Figure 6E). This fits with previous findings that the antioxidant capacity of FLO-1 is significantly lower than that of OAC-P4C ^37^. This inverse correlation of sensitivity between elesclomol and disulfiram and H_2_O_2_ suggests that elesclomol and disulfiram activity is unrelated to ROS, at least in OAC.

**Figure 6:**
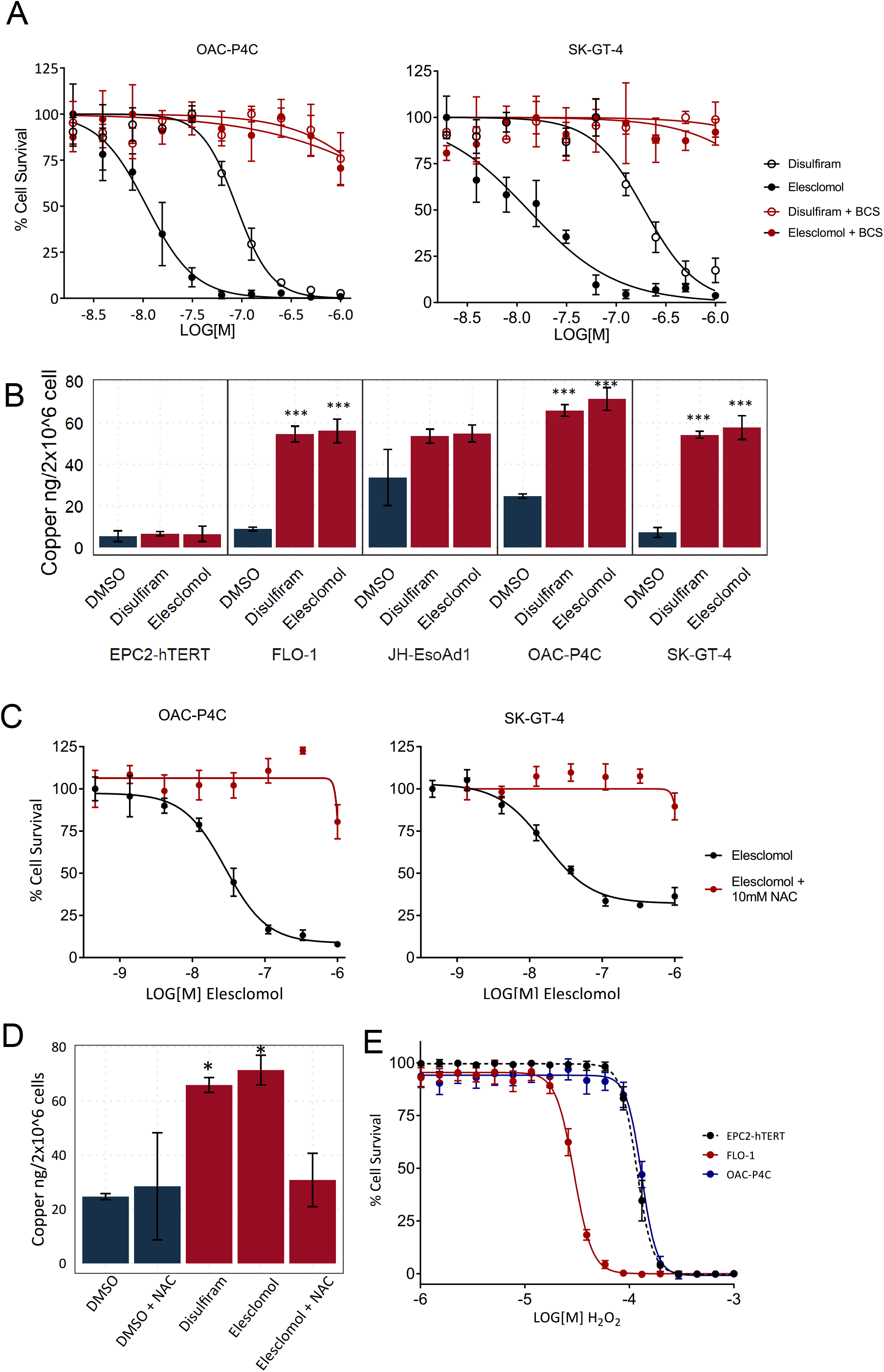
Mechanistic studies. A) Dose responses with and without copper chelator bathocuproinedisulfonic acid. B) ICP-MS intracellular copper levels. Error bars indicate SE. 1-ANOVA and Tukeys post hoc test for significance. Significance indicated for treatments compared to DMSO. *** p<0.001, n = 3. C) Elesclomol dose response with and without N-acetyl-L-cysteine (NAC) pre-treatment * P < 0.05. D) ICP-MS intracellular copper levels in the OAC cell line OAC-P4C with and without 30 min, 10 μM NAC pre-treatment. E) H_2_O_2_ dose responses for OAC cell lines OAC-P4C and FLO-1, and tissue matched control EPC2-hTERT.

### Copper is essential for compound activity

A recent microarray analysis of copper induced gene expression in a human lung epithelial cell line ^38^ identified the heat shock response, unfolded protein response, and ubiquitin/proteasomal degradation pathways. Further, it identified transcriptional downregulation of H_2_O_2_ sink enzymes, demonstrating remarkable similarity to the gene expression signature produced by disulfiram and elesclomol in OAC, suggesting copper may be involved in the mechanism of these three compounds, as has been proposed for elesclomol previously ^36^.

To interrogate whether copper plays a role in the activity of these compounds, we assessed whether copper was an essential component of the cell media. Pre-incubation of cells with cell impermeable bathocuproinedisulfonic acid to remove free copper from the media led to complete loss of activity for both disulfiram and elesclomol (Figure 6A), confirming free extracellular copper is necessary for the cytotoxic activity of both compounds. The iron chelators deferoxamine (data not shown) and ciclopirox olamine (Supplementary Figure 7) were unable to rescue cell death, demonstrating specificity for copper.

Using inductively coupled plasma mass spectrometry (ICP-MS) to detect metal ions within cell extracts we found that both disulfiram and elesclomol treatment led to the accumulation of intracellular copper in OAC cells (Figure 6B) identifying them as copper ionophores in OAC. We also found that other metal ions; Fe, Mg, Mn, and Zn, did not increase with compound treatment (Supplementary Figure 8 for Fe levels and data not shown for other metal ions), again demonstrating the specificity for copper in their mechanism of action. Consistent with the lack of gene expression changes, ICP-MS showed no accumulation of copper in the tissue-matched control cells after incubation with either disulfiram or elesclomol (Figure 6B).

N-acetyl-l-cysteine (NAC) is an antioxidant and has previously been shown to alleviate elesclomol’s activity ^39,40^ (Figure 6C), thus, elesclomol is thought to act via ROS induction. Importantly we demonstrate here for the first time that pre-incubation with the antioxidant NAC leads to complete loss of copper accumulation post elesclomol treatment, and results in copper levels comparable with those of the untreated cells (Figure 6D). Therefore NAC pre-incubation appears to prevent compound activity by preventing the increase in intracellular copper levels (Figure 6B) and thus is not exclusively linked to ROS induction.

### Comparison between copper ionophores and proteasome inhibitor activity

Given the identification of upregulated heat shock response and the unfolded protein response, known to arise from increased levels of misfolded proteins in the cytosol and endoplasmic reticulum respectively ^35^, and ubiquitin and proteasome subunit expression correlating with sensitivity, we went back to study potential links to protein misfolding and proteostasis in order to identify the mechanism by which these compounds work since it does not appear to be via ROS.

We quantified the phenotypic responses following treatment with proteasome inhibitors MG132 and Lactacystin (Figure 7, green points) and compared them with disulfiram, elesclomol, and ammonium pyrrolidinedithiocarbamate to determine if they produced a similar cellular phenotype, which would suggest proteasome inhibition may be their mechanism of action. However, phenotypically they do not cluster together, indicating that these copper ionophores do not induce a proteasome inhibitor like phenotype (Figure 7), indicating an alternative mechanism. However, when comparing the proteasome inhibitors to the copper ionophores, there is an inverse correlation of phenotypic activity across the cell lines. The proteasome inhibitors caused very limited changes in the OAC cell lines, with the exception of JH-EsoAD1, but caused strong phenotypic changes in the tissue-matched control cell lines, and the opposite is true for the copper ionophores (Figure 7) suggesting an inverse connection between the two groups of compounds. To further explore this association we compared the gene expression signatures induced by bortezomib, a proteasome inhibitor, and elesclomol (data from Tsvetkov et al., 2019) ^41^. Unexpectedly given the lack of similarity in phenotypic response, the two gene expression signatures show significant correlation (Pearson correlation 0.49, p< 0.001) (Figure 7D), although much lower than that between disulfiram and elesclomol. When taken together we suggest that this similarity in gene expression is because these copper ionophores elicit the same downstream response in cells as the proteasome inhibitors; dysregulation of proteostasis, but critically we believe this to be via a mechanism that is distinct from proteasome inhibition, given the dissimilarity in morphological signatures and the opposing sensitivity of cells to the two groups of compounds.

**Figure 7:**
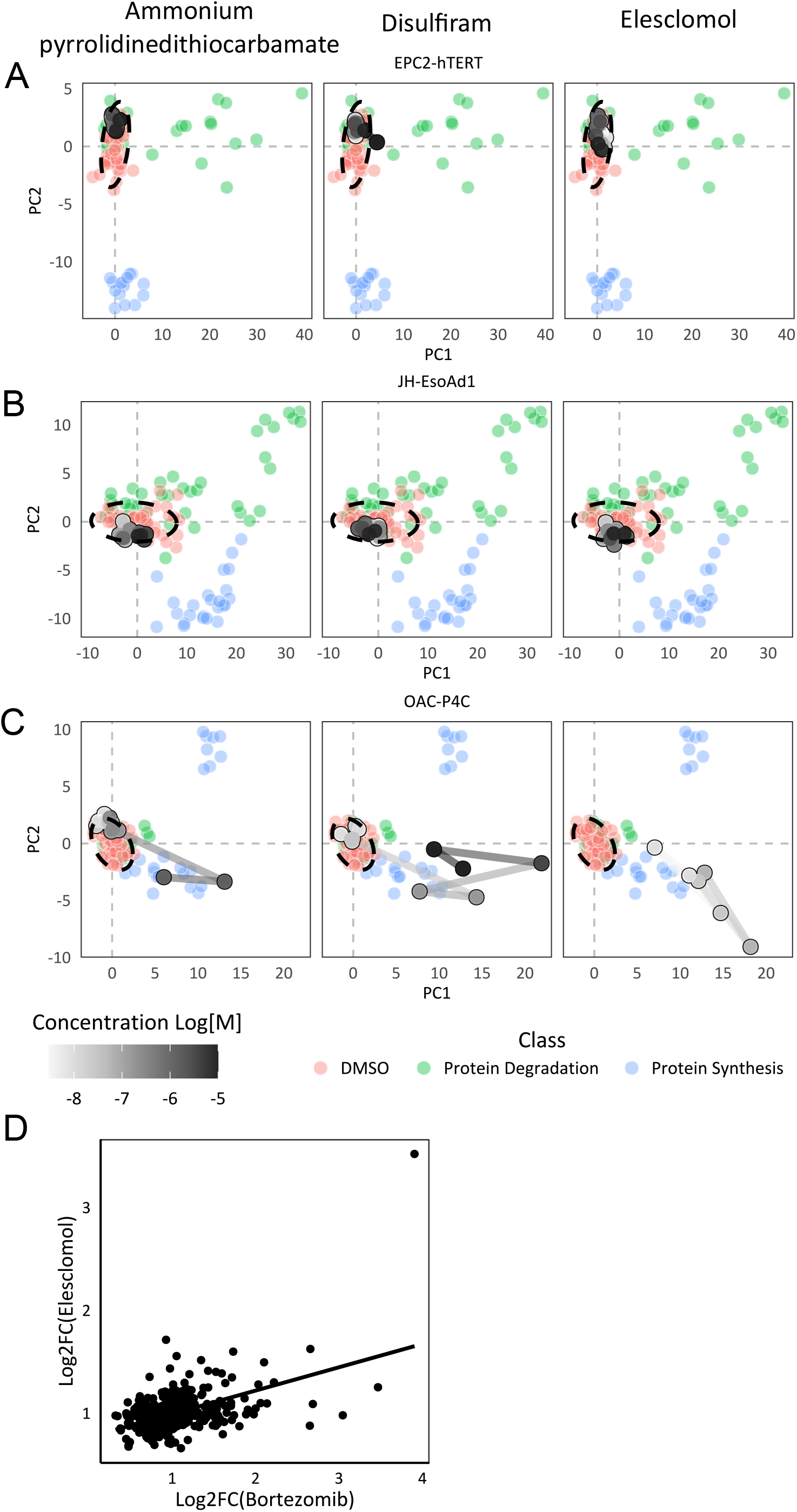
Comparison to compounds affecting proteostasis. First two principal components for A) Tissue matched cell line EPC2-hTERT. B) Elesclomol insensitive cell line JH-EsoAd1. C) Elesclomol sensitive cell line OAC-P4C. Reference library coloured my mechanistic class, dose response coloured by concentration. D) Correlation of bortezomib and elesclomol induced gene expression changes. (Data from Tsvetkov et al., 2019.)

Overall, our data indicate elesclomol, disulfiram and ammonium pyrrolidinedithiocarbamate act via a shared cancer-cell-specific mechanism involving an increase in cellular copper levels leading to dysregulation of proteostasis and selective cytotoxicity. Thus we present the identification of a new therapeutic mechanism for the treatment of OAC.

## Discussion

Elesclomol, disulfiram, and ammonium pyrrolidinedithiocarbamate have been linked to multiple and sometimes conflicting targets and modes of action; for example ammonium pyrrolidinedithiocarbamate demonstrates antioxidant capacity ^21^, while elesclomol is thought to induce ROS ^25,39,40,42^ and the mechanism-of-action of disulfiram has been attributed to several targets including NF-kappaB and the proteasome ^23^. In this work, systematic assessment of these compounds in OAC using a panel of cell lines displaying distinct sensitivities to the compounds has begun to unravel which of these mechanisms are relevant to their anticancer activity at physiologically relevant doses. Firstly, comparison of compound activity across our panel of cell lines and correlation between the morphological and gene expression signatures of the compounds revealed a shared mechanism of action between all three compounds not previously described in the literature. In pursuit of identification of the mechanism responsible for their selective activity in OAC this work has confirmed that the mechanism is copper dependent and leads to the accumulation of intracellular copper. We believe that all cellular responses to these compounds are likely entirely due to the accumulation of copper and the compounds act as a shuttle, bypassing homeostatic copper regulatory mechanisms within a cell. This would explain why the induced gene expression signature is so similar to that of the very high levels of extracellular copper in other published work ^38^. All three compounds share structural motifs which can lead to the chelation of copper outside of cells ^36,43–45^. However, the mechanism by which these compounds lead to the accumulation of copper within OAC cells specifically is still unknown and warrants further investigation as this may prove to be a novel strategy for treating OAC.

It has previously been thought that elesclomol worked via induction of ROS due to alleviation of its effects by NAC ^36,39,46^. However, work carried out here demonstrates that NAC is actually interfering upstream of ROS induction, preventing the accumulation of intracellular copper. One possible explanation is that NAC has been shown to potently reduce Cu(II) to Cu(I) ^38^ and this may prevent the elesclomol-copper chelate formation, while disulfiram requires Cu(II) for its conversion to an active metabolite diethylthiolcarbamate (DETC) ^43^ so in the presence of Cu(I) it will not undergo conversion to form copper chelates, explaining its lack of activity. We also find that the most sensitive cell line to the copper ionophores (OAC-P4C) is more resistant to oxidative stress induction by H_2_O_2_ treatment than the elesclomol insensitive cell line FLO-1, and the compounds do not induce expression of known antioxidant enzymes, further suggesting the mechanism is not related to ROS.

Gene expression studies revealed strong induction of the heat shock response, the NRF2-mediated oxidative stress response, unfolded protein response, and endoplasmic reticulum stress, pointing towards dysregulation of proteostasis as the mechanism through which the compounds act. Utilising proteasome inhibitors we made several unique predictions from the morphological data. Firstly we identified an inverse correlation in morphological activity between proteasome inhibitors and our copper ionophores which we correctly hypothesised and indicates that proteasome inhibitor resistant cells display a unique sensitivity to the copper ionophores, as has now been proven for elesclomol using proteasome inhibitor resistant cells ^47^. Secondly, given the dissimilarity in morphological signatures we propose that the copper ionophores act via a mechanism that is distinct from the proteasome inhibitors. Crucially though, from the gene expression similarity between elesclomol and bortezomib, a proteasome inhibitor, we believe that the copper ionophores still elicit the same downstream response in cells as the proteasome inhibitors; dysregulation of proteostasis. This fits with the induced gene expression signature. We provide further evidence towards a mechanism involving dysregulation of proteostasis through the identification that proteasome pathway basal gene expression levels predict sensitivity to elesclomol and disulfiram from GSEA analysis. This proposed mechanism also explains why elesclomol was recently found to synergise with the proteasome inhibitor bortezomib ^47^, since we believe them to both cooperate and reinforce the same downstream consequence in cells.

With regards to the sensitivity of OAC cell lines to elesclomol, disulfiram and ammonium pyrrolodinedithiocarbamate, this work has identified differences in metabolism and proteostasis that we believe contribute to copper ionophore sensitivity in OAC. Given the recently identified associations between mitochondrial function and the ubiquitin-proteasome system ^47–50^, unravelling the interplay between the two will lead to a greater understanding of both proteasome inhibitor and copper ionophore resistance and how the two can be used in combination. Further we identify a number of potential genes which may define sensitivity. At the single gene level and using GSEA, both OAC specific and pan cancer analyses revealed methionine sulfoxide reductase (MSR) genes and pathways as potential biomarkers of sensitivity. MSR genes have been identified as upregulated in cancer previously, for example MSRB3 is significantly upregulated in clear cell renal carcinoma (RCC) samples compared to adjacent tissues ^51^. In line with these findings, it has been shown that copper affects the stability and/or assembly of iron-sulfur cluster proteins in eukaryotic cells ^52^, and that knockout of the MSR genes in yeast leads to a copper resistance phenotype and is related to iron-sulfur cluster function ^53^. These previous studies demonstrate a relationship between copper sensitivity and MSR gene expression in yeast and other organisms. Our finding here suggest that this extends to elesclomol and copper ionophore induced copper toxicity in human OAC, and it could be used as a biomarker for sensitivity.

In summary, this work provides unique examples of how high-content profiling data in OAC can be utilised to identify, characterise and study associations in compound activity across panels of distinct cell lines to provide insights into new therapeutic approaches, cancer vulnerabilities, and patient stratification hypotheses to advance drug discovery and form the basis for future biomarker-based clinical trials in OAC.

