## Supplementary Tables and Figures for "High-content profiling reveals a unified model of copper ionophore dependent cell death in oesophageal adenocarcinoma"

Supplementary Table 1: Compound libraries. CRUK therapeutics discovery laboratories Library (CRT). Library of pharmacologically active compounds (LOPAC).

| Library | Number of Plates | Source Plate Concentration (µM) | Assay Concentration (µM) |
| --- | --- | --- | --- |
| Prestwick | 4 | 1,000 | 1 |
| LOPAC | 5 | 3,000 | 3 |
| Bespoke | 1 | 1,000 | 1 |
| BioAscent | 10 | 10,000 | 10 |
| CRT | 44 | 10,000- 12,000 | 10-12 |

Supplementary Table 2: Patient and tumour origin and characteristics of oesophageal adenocarcinoma cell lines. GOJ, gastroesohageal junction

| Cell Line | Gender | Age | Ethnicity | Barrett’s | Location | Grade | Stage | Sequencing study |
| --- | --- | --- | --- | --- | --- | --- | --- | --- |
| SK-GT-4 | Male | 89 | White | Yes | Distal 1/3 | Well | pT2N1Mx | ^1^ |
| FLO-1 | Male | 68 | White | No | Distal 1/3 | Poor | pT2N1M0 | ^1,2^ |
| OE33 | Female | 73 | White | Yes | Distal 1/3 | Poor | pT3N0M0 | ^1,2^ |
| OAC-P4C | Male | 55 | White | No | GOJ | Moderate | pT3N1M1 | ^1^ |
| JH-EsoAd1 | Male | 66 | White | Yes | Distal 1/3 | Moderate | pT3N0M0 | ^1^ |
| MFD-1 | Male | 55 | White | No | Distal 1/3 | Moderate | pT4N3M0 | ^2^ |

1.	Contino G, Eldridge MD, Secrier M, et al. Whole-genome sequencing of nine esophageal adenocarcinoma cell lines. F1000Research 2016;5:1336.

2.	Garcia E, Hayden A, Birts C, et al. Authentication and characterisation of a new oesophageal adenocarcinoma cell line: MFD-1. Sci Rep 2016;6:32417.

**Supplementary Table 3: Cell Painting reagents.** Concentrations, excitation/emission wavelengths of the filters used for imaging, and suppliers. ex: excitation, em: emission

| **Stain** | **Structure** | **Wavelength**  **(ex/em [nm])** | **Channel** | **Concentration** | **Cat No; Supplier** |
| --- | --- | --- | --- | --- | --- |
| Hoescht 33342 | Nuclei | 387/447 | DAPI | 4 µg/mL | #H1399; Mol. Probes |
| SYTO 14 | Nucleoli | 531/593 | CY3 | 3 µM | #S7576; Invitrogen |
| Phalloidin 594 | F-actin | 562/624 | TxRED | 0.14X | #ab176757; Abcam |
| Wheat germ agglutinin Alexa Fluor 594 | Golgi and Plasma Membrane | 562/624 | TxRED | 1 µg/mL | #W11262; Invitrogen |
| Concanavalin A Alexa Fluor 488 | Endoplas-mic Reticulum | 462/520 | FITC | 20 µg/mL | #C11252; Invitrogen |
| MitoTracker DeepRed | Mitochond-ria | 628/692 | CY5 | 600 nM | #M22426; Invitrogen |

**Table 4: Seahorse assay conditions.** FCCP- Trifluoromethoxy carbonylcyanide phenylhydrazone

| Cell Line | Seeding Density (per well) | Concentration (μM) | | |
| --- | --- | --- | --- | --- |
|  |  | Oligomycin | FCCP | Antimycin/ Rotenone |
| OAC-P4C | 10,000 | 2 | 0.6 | 3 |
| SK-GT-4 | 8,000 | 1 | 1.2 | 2 |
| FLO-1 | 8,000 | 1 | 1.2 | 3 |
| JH-EsoAd1 | 10,000 | 1 | 2.4 | 1 |

Supplementary Table 5: Primary screen compound hits. *Not available for rescreening

|  | Phenotypic hits | cell survival hits |
| --- | --- | --- |
| 1 | (S)-(+)-Camptothecin |  |
| 2 | 5-Fluorouracil | Adrucil(Fluorouracil) |
| 3 | Alcuronium chloride |  |
| 4 | Aminopterin |  |
| 5 |  | Ammonium pyrrolidinedithiocarbamate |
| 6 | Amonafide |  |
| 7 | Amsacrine hydrochloride |  |
| 8 | Ancitabine hydrochloride |  |
| 9 | Ascorbic acid | Ascorbic acid |
| 10 |  | Aurora A Inhibitor I |
| 11 | Azapropazone |  |
| 12 | AZD7762 |  |
| 13 | Bacampicillin hydrochloride* | Bacampicillin hydrochloride* |
| 14 | Benzylpenicillin sodium | Benzydamine hydrochloride |
| 15 | Betahistine mesylate |  |
| 16 | BI 2536 |  |
| 17 | BIX-01294·3HCl |  |
| 18 | Calcimycin |  |
| 19 | Cantharidic Acid |  |
| 20 | Cantharidin |  |
| 21 | Carmofur |  |
| 22 | CCT137690 |  |
| 23 | Ceforanide |  |
| 24 | CID 11210285 hydrochloride |  |
| 25 |  | Clindamycin hydrochloride |
| 26 | Clofarabine |  |
| 27 |  | Clorgyline hydrochloride |
| 28 | Colchicine |  |
| 29 | Cytosine-1-beta-D-arabinofuranoside hydrochloride |  |
| 30 | Deptropine citrate | Deptropine citrate |
| 31 | Dipivefrin hydrochloride | Dipivefrin hydrochloride |
| 32 |  | Disulfiram |
| 33 |  | Elesclomol |
| 34 | Ellence |  |
| 35 |  | Etifenin |
| 36 | Etoposide |  |
| 37 | Floxuridine | Floxuridine |
| 38 | Fluoxetine hydrochloride | Fluoxetine hydrochloride |
| 39 | Gemcitabine hydrochloride |  |
| 40 | Ibuprofen |  |
| 41 | JNJ-26481585 |  |
| 42 | Letrozole |  |
| 43 | Methotrexate | Methotrexate |
| 44 | Mitoxantrone |  |
| 45 | Mizolastine |  |
| 46 | MK-2206 | MK-2206 |
| 47 | MLN8237 |  |
| 48 | Molsidomine |  |
| 49 |  | Nicergoline |
| 50 | NSC-3852 | NSC-3852 |
| 51 | Obatoclax Mesylate |  |
| 52 |  | Oxytetracycline dihydrate |
| 53 | Paclitaxel |  |
| 54 | Pemetrexed disodium (LY-231514) | Pemetrexed disodium (LY-231514) |
| 55 | Pirenperone |  |
| 56 | PKC-412 |  |
| 57 | PMEG hydrate* |  |
| 58 | Podophyllotoxin |  |
| 59 |  | Propylthiouracil |
| 60 | Raltitrexed (Tomudex) | Raltitrexed(Tomudex) |
| 61 | SB743921 hydrochloride |  |
| 62 | SB939 |  |
| 63 | SNS-032(BMS-387032) |  |
| 64 | SNS-314 Mesylate |  |
| 65 | Suprafenacine |  |
| 66 |  | Terbutaline hemisulfate |
| 67 | Thiocolchicine |  |
| 68 | Thiorphan | Thiorphan |
| 69 | Topotecan hydrochloride hydrate |  |
| 70 | Trichostatin A | Trichostatin A |
| 71 | Vincristine sulfate |  |
| 72 | Vinorelbine (Navelbine) |  |
| 73 | VX-680 |  |
| 74 |  | YM155 |

Supplementary Table 6. Dose response validation compounds. (Red- not selective)

|  | validation compounds | Phenotypic Validation | Cell survival Validation |
| --- | --- | --- | --- |
| 1 | (S)-(+)-Camptothecin | ✓ | ✓ |
| 2 | 5-Fluorouracil | ✓ | ✓ |
| 3 | alcuronium chloride |  |  |
| 4 | Aminopterin |  | ✓ |
| 5 | Ammonium pyrrolidinedithiocarbamate | ✓ | ✓ |
| 6 | Amonafide | ✓ | ✓ |
| 7 | Amsacrine hydrochloride | ✓ | ✓ |
| 8 | Ancitabine hydrochloride | ✓ | ✓ |
| 9 | Ascorbic acid |  |  |
| 10 | Aurora A Inhibitor I | ✓ | ✓ |
| 11 | Azapropazone |  |  |
| 12 | AZD7762 | ✓ | ✓ |
| 13 | Benzydamine hydrochloride | ✓ |  |
| 14 | Benzylpenicillin sodium |  |  |
| 15 | Betahistine mesylate |  |  |
| 16 | BI 2536 | ✓ | ✓ |
| 17 | BIX 01294 trihydrochloride hydrate | ✓ |  |
| 18 | Calcimycin | ✓ | ✓ |
| 19 | Cantharidic Acid | ✓ |  |
| 20 | Cantharidin | ✓ | ✓ |
| 21 | Carmofur | ✓ | ✓ |
| 22 | CCT137690 | ✓ | ✓ |
| 23 | Ceforanide |  |  |
| 24 | CID 11210285 hydrochloride | ✓ | ✓ |
| 25 | Clindamycin hydrochloride |  |  |
| 26 | Clofarabine | ✓ | ✓ |
| 27 | Clorgyline hydrochloride |  |  |
| 28 | Colchicine | ✓ | ✓ |
| 29 | Cytosine-1-beta-D-arabinofuranoside hydrochloride | ✓ | ✓ |
| 30 | deptropine citrate |  |  |
| 31 | Dipivefrin hydrochloride |  |  |
| 32 | Disulfiram | ✓ | ✓ |
| 33 | Elesclomol | ✓ | ✓ |
| 34 | Ellence | ✓ | ✓ |
| 35 | Etifenin |  |  |
| 36 | Etoposide | ✓ | ✓ |
| 37 | Floxuridine | ✓ | ✓ |
| 38 | Fluoxetine hydrochloride | ✓ |  |
| 39 | Gemcitabine hydrochloride | ✓ | ✓ |
| 40 | Ibuprofen (racemic) |  |  |
| 41 | JNJ-26481585 | ✓ | ✓ |
| 42 | Letrozole |  |  |
| 43 | Methotrexate | ✓ | ✓ |
| 44 | Mitoxantrone | ✓ | ✓ |
| 45 | Mizolastine |  |  |
| 46 | MK-2206 | ✓ | ✓ |
| 47 | MLN8237 | ✓ | ✓ |
| 48 | Molsidomine |  |  |
| 49 | Nicergoline |  |  |
| 50 | NSC-3852 | ✓ | ✓ |
| 51 | Obatoclax Mesylate | ✓ | ✓ |
| 52 | Oxytetracycline dihydrate |  |  |
| 53 | Paclitaxel | ✓ | ✓ |
| 54 | Pemetrexed disodium (LY-231514) | ✓ | ✓ |
| 55 | pirenperone |  |  |
| 56 | PKC-412 |  | ✓ |
| 57 | Podophyllotoxin |  | ✓ |
| 58 | Propylthiouracil |  |  |
| 69 | Raltitrexed(Tomudex) | ✓ | ✓ |
| 60 | SB743921 hydrochloride | ✓ | ✓ |
| 61 | SNS-032(BMS-387032) | ✓ | ✓ |
| 62 | SNS-314 (mesylate) | ✓ | ✓ |
| 63 | Suprafenacine | ✓ | ✓ |
| 64 | Terbutaline hemisulfate |  |  |
| 65 | Thiocolchicine | ✓ | ✓ |
| 66 | Thiorphan |  |  |
| 67 | Topotecan hydrochloride hydrate | ✓ | ✓ |
| 68 | Trichostatin A | ✓ | ✓ |
| 69 | Vincristine sulfate | ✓ | ✓ |
| 70 | Vinorelbine(Navelbine) | ✓ | ✓ |
| 71 | VX-680 | ✓ | ✓ |
| 72 | YM155 | ✓ | ✓ |

Supplementary Table 7: Compound IC_50_ values across panel of cell lines.

|  | IC_50_ (nM) | | |
| --- | --- | --- | --- |
| Cell Line | Elesclomol | Disulfiram | Ammonium pyrrolidinedithiocarbamate |
| CP-A | >10,000 | >10,000 | >10,000 |
| EPC2-hTERT | >10,000 | >10,000 | >10,000 |
| FLO-1 | >10,000 | >10,000 | >10,000 |
| JH-EsoAd1 | 3582 | >10,000 | >10,000 |
| MFD-1 | 51 | 868 | 1463 |
| OE33 | 39 | >10,000 | 1317 |
| SK-GT-4 | 5 | 197 | 915 |
| OAC-P4C | 1 | 69 | 880 |

Supplementary Table 8: Patient derived organoids. Elesclomol IC_50_ values.

|  |  | **IC_50_** | |
| --- | --- | --- | --- |
| Cell Line | Type | Log [M] | nM |
| NG088 | Normal Gastric | -5.36 | 4325 |
| CAM401 | OAC | -6.83 | 147 |
| CAM408 | OAC | -7.65 | 22 |
| CAM277 | OAC | -7.98 | 10 |
| CAM468 | OAC | -8.25 | 5.6 |
| CAM479 | OAC | -9.77 | 0.17 |

Supplementary Table 9: Mechanism of action reference compounds. Compound name, mechanism of action, sub-class and supplier and catalogue number.

| **Compound Name** | **Mechanism of Action** | **Sub-Class** | **Supplier; Catalogue number** |
| --- | --- | --- | --- |
| Cytochalasin B | Actin disrupting | Actin disruptor | Sigma; C8273 |
| Cytochalasin D | Actin disrupting | Actin disruptor | Sigma; C6762 |
| Latrunculin | Actin disrupting | Actin stabiliser | Sigma; L5288 |
| Camptothecin | DNA damaging | Topoisomerase-1 inhibitor | Selleckchem; S1288 |
| SN38 | DNA damaging | Topoisomerase-1 inhibitor | Selleckchem; S4908 |
| Dasatinib | Kinase inhibitor | Src- EMT | Selleckchem; S1021 |
| Saracatinib | Kinase inhibitor | Src-EMT | Selleckchem; S1006 |
| Epothilone B | Microtubule disrupting | Microtubule stabiliser | Selleckchem; S1364 |
| Paclitaxel | Microtubule disrupting | Microtubule stabiliser | Sigma; T7402 |
| Colchicine | Microtubule disrupting | Microtubule destabiliser | Sigma; C9754 |
| Nocodazole | Microtubule disrupting | Microtubule destabiliser | Sigma; M1404 |
| Monastrol | Microtubule disrupting | Eg5 kinesin inhibitor | Sigma; M8515 |
| ARQ621 | Microtubule disrupting | Eg5 kinesin inhibitor | Selleckchem; S7355 |
| Barasertib | Microtubule disrupting | Aurora kinase B inhibitor | Selleckchem; S1147 |
| ZM447439 | Microtubule disrupting | Aurora kinase B inhibitor | Selleckchem; 1103 |
| MG132 | Protein degradation | Proteasome | Selleckchem; S2619 |
| Lactacystin | Protein degradation | Proteasome | Tocris; 2267 |
| ALLN | Protein degradation | Cysteine/calpain | Sigma; A6165 |
| ALLM | Protein degradation | Cysteine/calpain | Sigma; A6060 |
| Cycloheximide | Protein synthesis | Protein synthesis | Sigma; 01810 |
| Emetine | Protein synthesis | Protein synthesis | Sigma; E2375 |
| Lovastatin | Statin | Statin | Sigma; PHR1285 |
| Simvastatin | Statin | Statin | Sigma; PHR1438 |
| SAHA | HDAC inhibitor | HDAC inhibitor | Sigma; SML0061 |
| Panobinostat | HDAC inhibitor | HDAC inhibitor | Selleckchem; S1030 |
| Trichostatin A | HDAC inhibitor | HDAC inhibitor | Selleckchem; S1045 |
| Romidepsin | HDAC inhibitor | HDAC inhibitor | Selleckchem; S3020 |
| Entinostat | HDAC inhibitor | HDAC inhibitor | Selleckchem; S1053 |
| Quisinostat | HDAC inhibitor | HDAC inhibitor | Selleckchem; S1096 |
| Ricolinostat | HDAC inhibitor | HDAC inhibitor | Selleckchem; S8001 |
| Tubastatin A | HDAC inhibitor | HDAC inhibitor | Selleckchem; S8049 |
| Droxinostat | HDAC inhibitor | HDAC inhibitor | Selleckchem; S1422 |
| PCI34051 | HDAC inhibitor | HDAC inhibitor | Selleckchem; S2021 |
| TMP195 | HDAC inhibitor | HDAC inhibitor | Selleckchem; S8502 |
| LMK235 | HDAC inhibitor | HDAC inhibitor | Selleckchem; S7569 |
| CUDC-907 | HDAC inhibitor | HDAC inhibitor | Selleckchem; S2759 |
| Belinostat | HDAC inhibitor | HDAC inhibitor | Selleckchem; S1085 |

Supplementary Table 10: Rank order for core enriched genes in the MYC 1 Hallmark Geneset.

| Number | Gene Symbol | Gene Name | Rank |
| --- | --- | --- | --- |
| 1 | UBA2 | ubiquitin like modifier activating enzyme 2 | 87 |
| 2 | PSMD8 | proteasome 26S subunit, non-ATPase 8 | 212 |
| 3 | SRM | spermidine synthase | 215 |
| 4 | LSM2 | LSM2 homolog, U6 small nuclear RNA and mRNA degradation associated | 267 |
| 5 | RACK1 | receptor for activated C kinase 1 | 287 |
| 6 | PSMC4 | proteasome 26S subunit, ATPase 4 | 375 |
| 7 | PWP1 | PWP1 homolog, endonuclein | 376 |
| 8 | UBE2E1 | ubiquitin conjugating enzyme E2 E1 | 479 |
| 9 | SF3A1 | splicing factor 3a subunit 1 | 507 |
| 10 | SF3B3 | splicing factor 3b subunit 3 | 575 |
| 11 | PABPC4 | poly(A) binding protein cytoplasmic 4 | 725 |
| 12 | PHB2 | prohibitin 2 | 751 |
| 13 | SLC25A3 | solute carrier family 25 member 3 | 860 |
| 14 | UBE2L3 | ubiquitin conjugating enzyme E2 L3 | 875 |
| 15 | USP1 | ubiquitin specific peptidase 1 | 887 |
| 16 | VDAC3 | voltage dependent anion channel 3 | 954 |
| 17 | SNRPA | small nuclear ribonucleoprotein polypeptide A | 1055 |
| 18 | XPOT | exportin for tRNA | 1133 |
| 19 | MRPL9 | mitochondrial ribosomal protein L9 | 1155 |
| 20 | SMARCC1 | SWI/SNF related, matrix associated, actin dependent regulator of chromatin subfamily c member 1 | 1161 |
| 21 | NAP1L1 | nucleosome assembly protein 1 like 1 | 1171 |
| 22 | RANBP1 | RAN binding protein 1 | 1353 |
| 23 | G3BP1 | G3BP stress granule assembly factor 1 | 1366 |
| 24 | PTGES3 | prostaglandin E synthase 3 | 1381 |
| 25 | HNRNPC | heterogeneous nuclear ribonucleoprotein C | 1384 |
| 26 | MRPS18B | mitochondrial ribosomal protein S18B | 1403 |
| 27 | CCT2 | chaperonin containing TCP1 subunit 2 | 1429 |
| 28 | SRPK1 | SRSF protein kinase 1 | 1490 |
| 29 | NOLC1 | nucleolar and coiled-body phosphoprotein 1 | 1512 |
| 30 | RAN | RAN, member RAS oncogene family | 1551 |
| 31 | HNRNPR | heterogeneous nuclear ribonucleoprotein R | 1565 |
| 32 | MCM5 | minichromosome maintenance complex component 5 | 1610 |
| 33 | CDK4 | cyclin dependent kinase 4 | 1636 |
| 34 | LSM7 | LSM7 homolog, U6 small nuclear RNA and mRNA degradation associated | 1652 |
| 35 | VBP1 | VHL binding protein 1 | 1661 |
| 36 | GNL3 | G protein nucleolar 3 | 1724 |
| 37 | CDC45 | cell division cycle 45 | 1769 |
| 38 | CSTF2 | cleavage stimulation factor subunit 2 | 1802 |
| 39 | CDC20 | cell division cycle 20 | 1807 |
| 40 | RSL1D1 | ribosomal L1 domain containing 1 | 1820 |
| 41 | KARS1 | lysyl-tRNA synthetase 1 | 1885 |
| 42 | U2AF1 | U2 small nuclear RNA auxiliary factor 1 | 1925 |
| 43 | PSMA4 | proteasome 20S subunit alpha 4 | 1931 |
| 44 | EIF3B | eukaryotic translation initiation factor 3 subunit B | 1978 |
| 45 | CAD | carbamoyl-phosphate synthetase 2, aspartate transcarbamylase, and dihydroorotase | 2065 |
| 46 | YWHAE | tyrosine 3-monooxygenase/tryptophan 5-monooxygenase activation protein epsilon | 2083 |
| 47 | FBL | fibrillarin | 2138 |
| 48 | TRA2B | transformer 2 beta homolog | 2140 |
| 49 | XPO1 | exportin 1 | 2151 |
| 50 | PPM1G | protein phosphatase, Mg2+/Mn2+ dependent 1G | 2423 |
| 51 | ETF1 | eukaryotic translation termination factor 1 | 2500 |
| 52 | TXNL4A | thioredoxin like 4A | 2578 |
| 53 | EIF4E | eukaryotic translation initiation factor 4E | 2646 |
| 54 | POLD2 | DNA polymerase delta 2, accessory subunit | 2664 |
| 55 | EIF3J | eukaryotic translation initiation factor 3 subunit J | 2667 |
| 56 | TUFM | Tu translation elongation factor, mitochondrial | 2700 |
| 57 | IMPDH2 | inosine monophosphate dehydrogenase 2 | 2734 |
| 58 | PSMB2 | proteasome 20S subunit beta 2 | 2808 |
| 59 | PSMD14 | proteasome 26S subunit, non-ATPase 14 | 2847 |
| 60 | EIF1AX | eukaryotic translation initiation factor 1A X-linked | 2855 |
| 61 | NPM1 | nucleophosmin 1 | 2877 |
| 62 | C1QBP | complement C1q binding protein | 2934 |
| 63 | BUB3 | BUB3 mitotic checkpoint protein | 3008 |
| 64 | POLE3 | DNA polymerase epsilon 3, accessory subunit | 3093 |
| 65 | RRP9 | ribosomal RNA processing 9, U3 small nucleolar RNA binding protein | 3205 |
| 66 | NHP2 | NHP2 ribonucleoprotein | 3209 |
| 67 | LDHA | lactate dehydrogenase A | 3235 |
| 68 | NOP16 | NOP16 nucleolar protein | 3266 |
| 69 | SNRPD3 | small nuclear ribonucleoprotein D3 polypeptide | 3334 |
| 70 | CCT4 | chaperonin containing TCP1 subunit 4 | 3422 |
| 71 | DEK | DEK proto-oncogene | 3531 |
| 72 | PA2G4 | proliferation-associated 2G4 | 3591 |
| 73 | CCT7 | chaperonin containing TCP1 subunit 7 | 3647 |
| 74 | NDUFAB1 | NADH:ubiquinone oxidoreductase subunit AB1 | 3668 |
| 75 | TCP1 | t-complex 1 | 3695 |
| 76 | PSMD7 | proteasome 26S subunit, non-ATPase 7 | 3716 |
| 77 | GLO1 | glyoxalase I | 3749 |
| 78 | CTPS1 | CTP synthase 1 | 3838 |
| 79 | PSMD1 | proteasome 26S subunit, non-ATPase 1 | 3883 |
| 80 | SSB | small RNA binding exonuclease protection factor La | 3961 |
| 81 | PCBP1 | poly(rC) binding protein 1 | 4034 |
| 82 | RPL14 | ribosomal protein L14 | 4035 |
| 83 | HPRT1 | hypoxanthine phosphoribosyltransferase 1 | 4109 |
| 84 | EPRS1 | glutamyl-prolyl-tRNA synthetase 1 | 4111 |
| 85 | TARDBP | TAR DNA binding protein | 4122 |
| 86 | EIF3D | eukaryotic translation initiation factor 3 subunit D | 4147 |
| 87 | MRPL23 | mitochondrial ribosomal protein L23 | 4192 |
| 88 | ODC1 | ornithine decarboxylase 1 | 4203 |
| 89 | RFC4 | replication factor C subunit 4 | 4218 |
| 90 | AIMP2 | aminoacyl tRNA synthetase complex interacting multifunctional protein 2 | 4249 |
| 91 | TYMS | thymidylate synthetase | 4371 |
| 92 | CYC1 | cytochrome c1 | 4420 |
| 93 | PRPS2 | phosphoribosyl pyrophosphate synthetase 2 | 4436 |
| 94 | APEX1 | apurinic/apyrimidinic endodeoxyribonuclease 1 | 4457 |

Supplementary Figures.

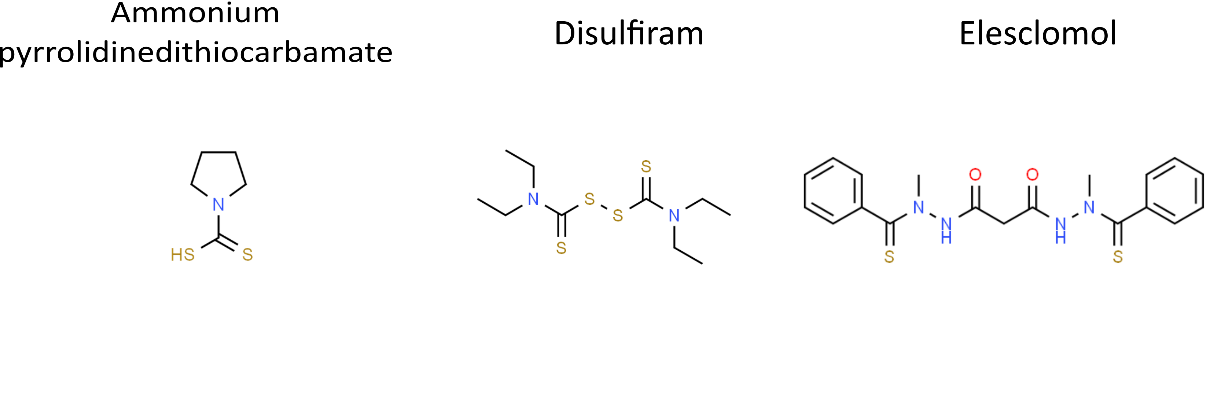

**Supplementary Figure 1: Chemical Structures of elesclomol, disulfiram and ammonium pyrrolidinedithiocarbamate.**

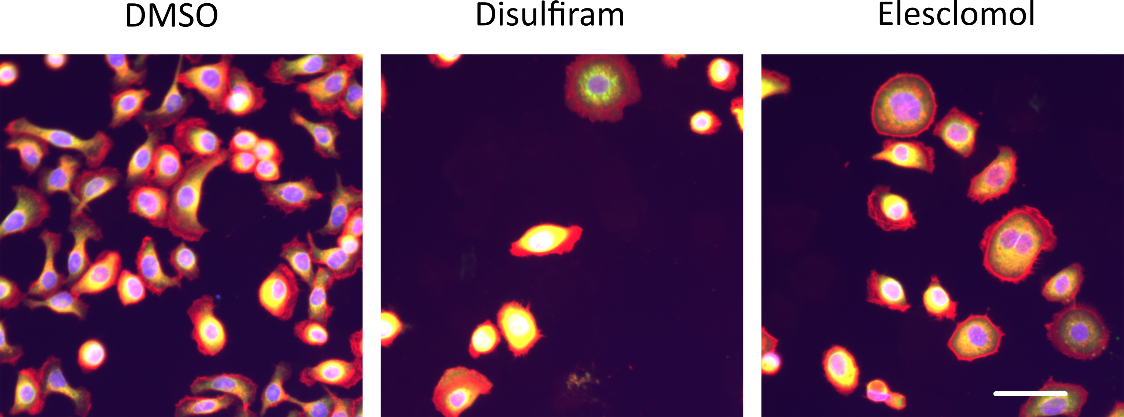

**Supplementary Figure 2: Colour combined images for DMSO, disulfiram (100 nM) and elesclomol (3 nM) treatments in the OAC-P4C cell line**. Hoescht 33342-nuclei (blue); Phalloidin 594, Wheat germ agglutinin and Alexa Fluor 594- F-actin, golgi and plasma membrane (red); Concanavalin A Alexa Fluor 488- Endoplasmic reticulum (green). Scale bars 50 µm.

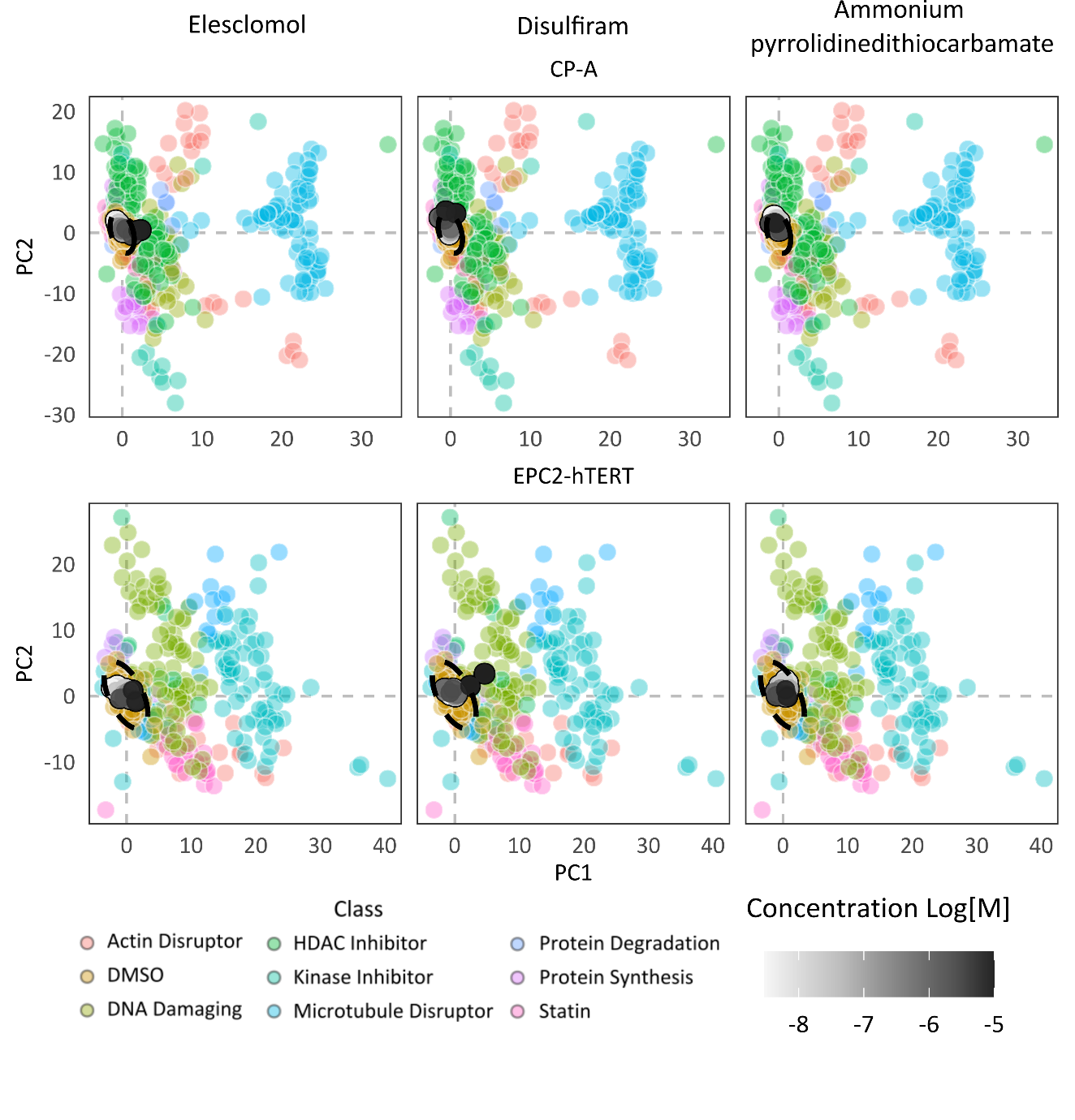

**Supplementary Figure 3: Phenotypic dose responses, for elesclomol, disulfiram and ammonium pyrrolidinedithiocarbamate in the tissue-matched control cell lines.** Library of reference compounds coloured by mechanistic class. Compound dose response overlay coloured by concentration.

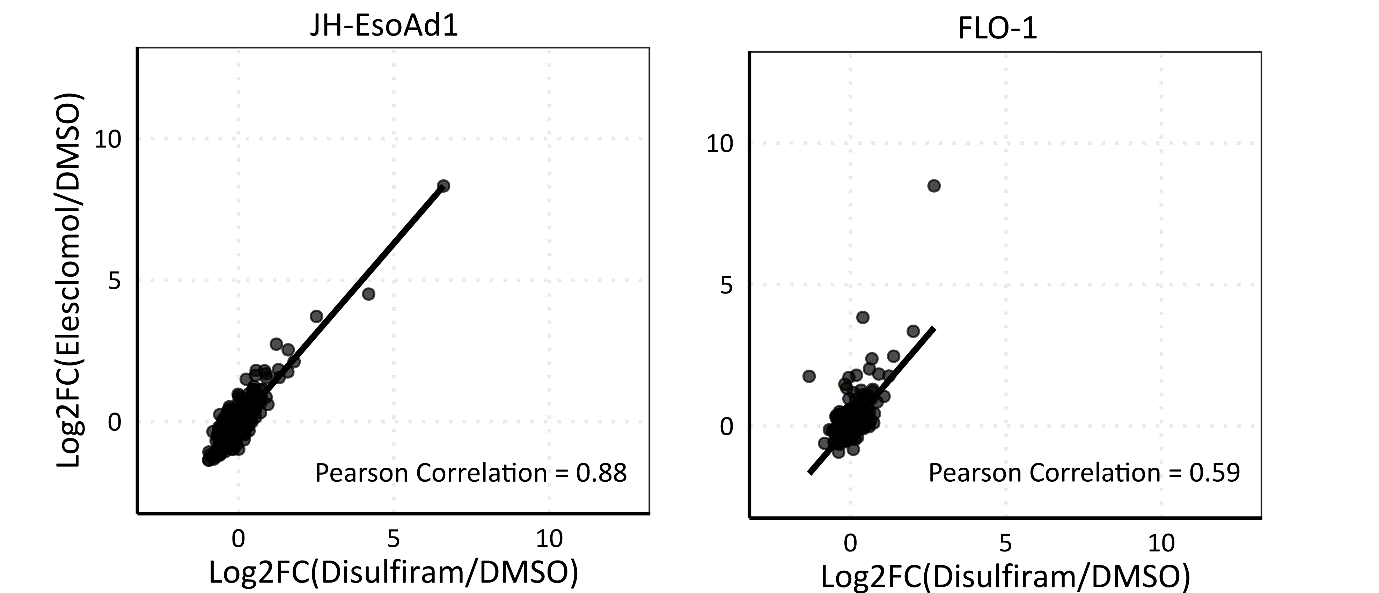

**Supplementary Figure 4: Disulfiram and Elesclomol induced log2 fold change gene expression changes in JH-EsoAd1 and FLO-1 cell lines.**

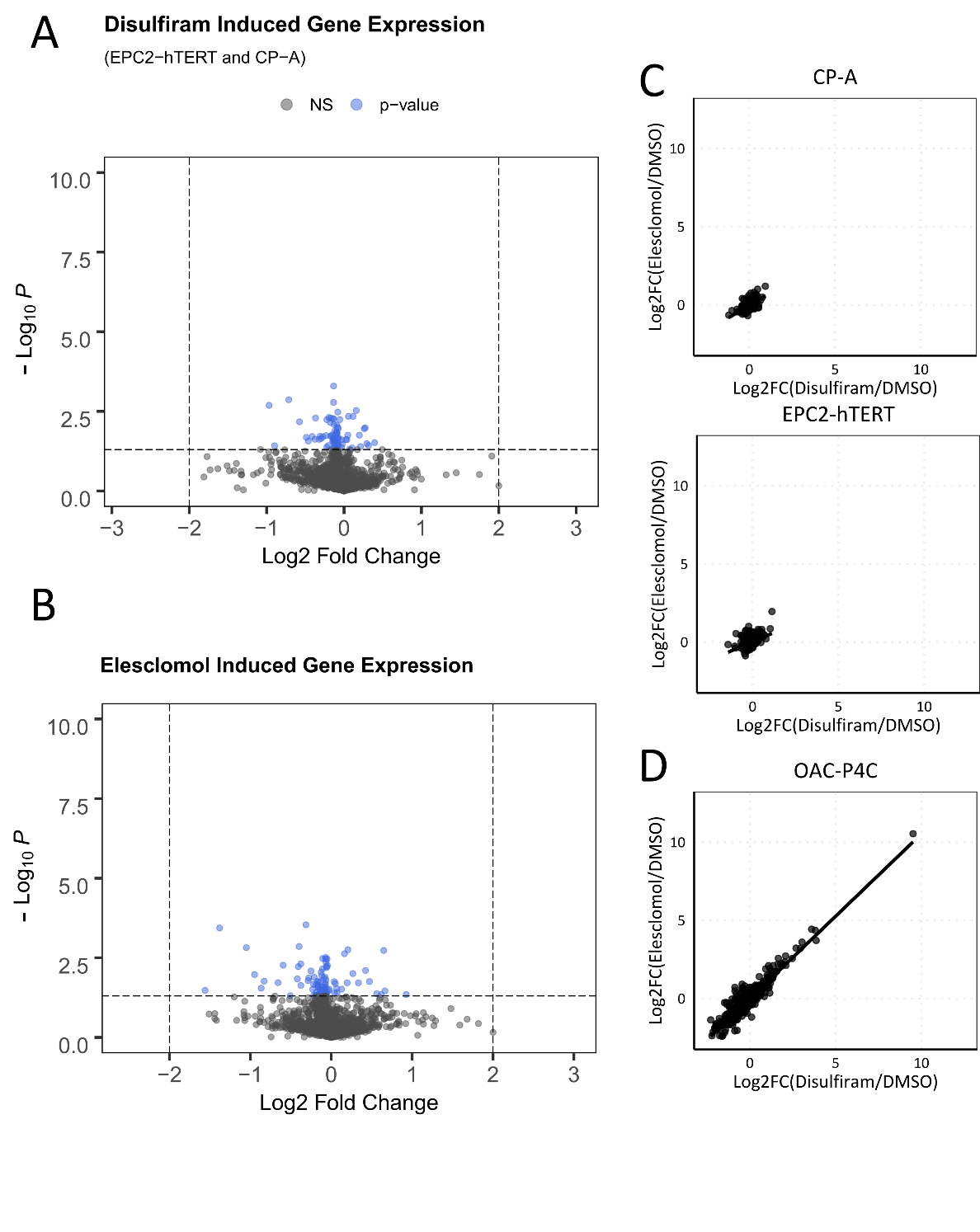

**Supplementary Figure 5: Treatment induced gene expression studies in tissue-matched control cell lines.** Differential expression analysis for A) Disulfiram, B) Elesclomol, induced gene expression for EPC2-hTERT and CP-A. n = 1. No significant genes after correcting for multiple testing. Correlation plot for Disulfiram and Elesclomol induced log2 fold change gene expression changes in C) tissue-matched controls CP-A and EPC2-hTERT, D) Sensitive OAC cell line OAC-P4C for comparison.

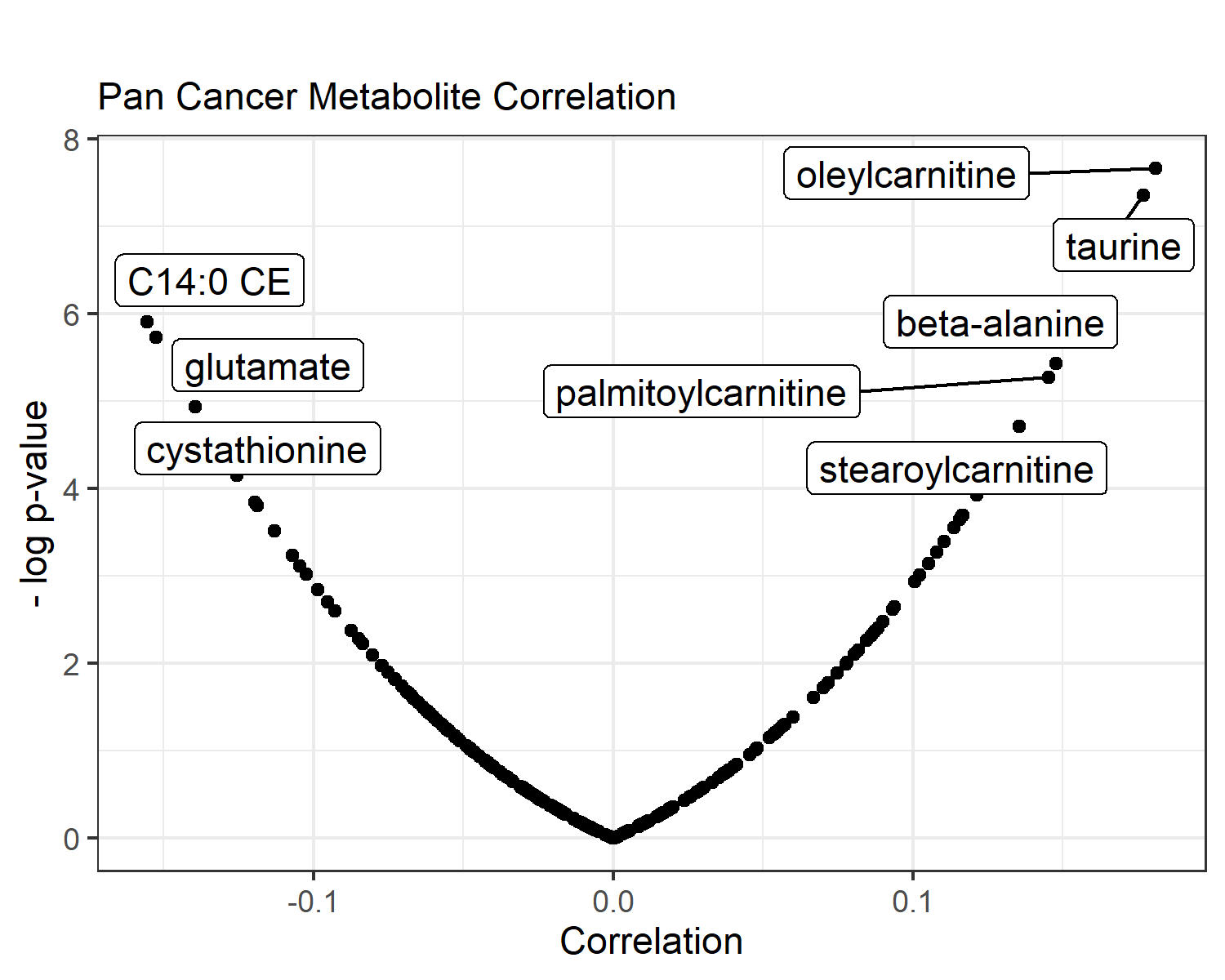

**Supplementary Figure 6: Pan Cancer Metabolite Correlation.**

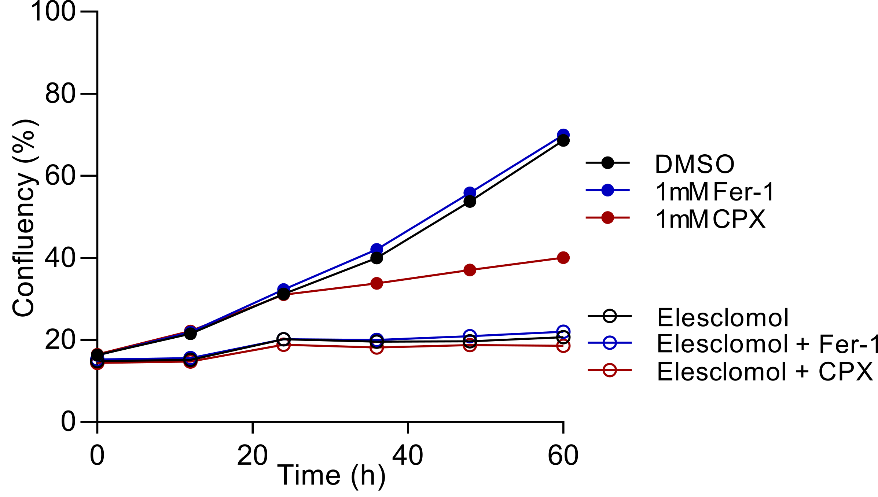

**Supplementary Figure 7: Role of iron.** OAC-P4C cell confluency after treatment with elesclomol (10 μM) in the presence or absence of the iron chelator ciclopirox olamine (CPX), and ferroptosis inhibitor Ferrostatin-1 (Fer-1).

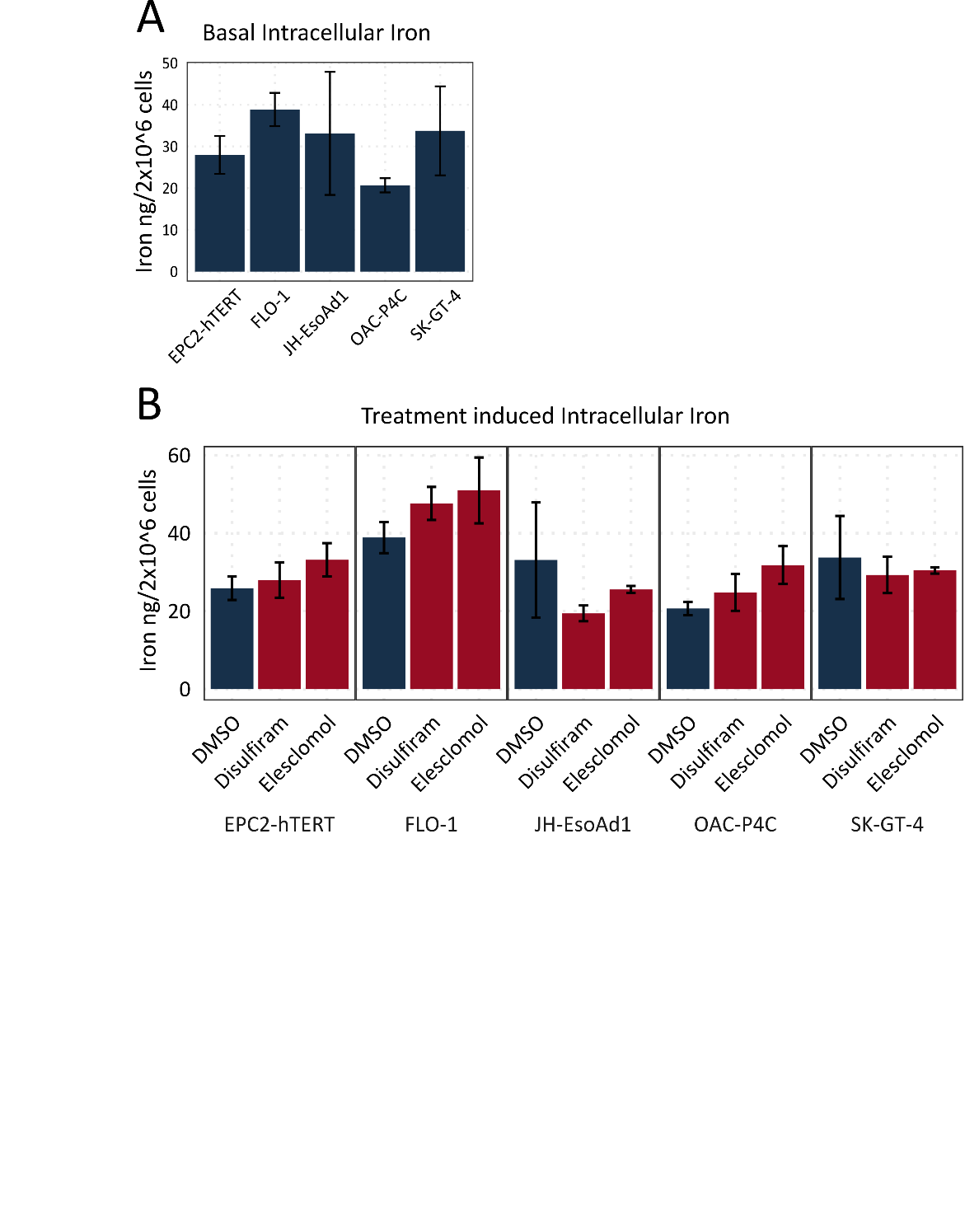

**Supplementary Figure 8: Iron ICP-MS.** A) Basal and B) Treatment induced intracellular iron levels determined by inductively coupled plasma mass spectrometry. Error bars indicate SE. n= 3.
